# Prediction of Transport, Deposition, and Resultant Immune Response of Nasal Spray Vaccine Droplets using a CFPD-HCD Model in a 6-Year-Old Upper Airway Geometry to Potentially Prevent COVID-19

**DOI:** 10.1101/2022.11.08.515673

**Authors:** Hamideh Hayati, Yu Feng, Xiaole Chen, Emily Kolewe, Catherine Fromen

## Abstract

This study focuses on the transport, deposition, and triggered immune response of intranasal vaccine droplets to the Angiotensin-converting enzyme 2-rich region (i.e., the olfactory region (OR)) in the nasal cavity of a 6-year-old female to possibly prevent COVID-19. To investigate how administration strategy can influence nasal vaccine efficiency, a validated multiscale model (i.e., computational fluid-particle dynamics (CFPD) and host-cell dynamics (HCD) model) was employed. Droplet deposition fraction, size change, residence time, and the area percentage of OR covered by the vaccine droplets and triggered immune system response were predicted with different spray cone angles, initial droplet velocities, and compositions. Numerical results indicate that droplet initial velocity and composition have negligible influences on the vaccine delivery efficiency to OR. In contrast, the spray cone angle can significantly impact the vaccine delivery efficiency. The triggered immunity was not significantly influenced by the administration investigated in this study, due to the low percentage of OR area covered by the droplets. To enhance the effectiveness of the intranasal vaccine to prevent COVID-19 infection, it is necessary to optimize the vaccine formulation and administration strategy so that the vaccine droplets can cover more epithelial cells in OR to minimize the available receptors for SARS-CoV-2.

## 4 Introduction

Intranasal vaccination against coronavirus disease 19 (COVID-19) caused by SARS-CoV-2 could be highly advantageous over conventional intramuscular vaccination. Indeed, the nasal cavity is the initial site of viral infection, replication, and transmission (Chen, Shen et al. 2020, Chavda, Vora et al. 2021, Pilicheva and Boyuklieva, Xi, Lei et al. 2021). After virus-laden droplets enter the nasal cavity, the spike (S) protein on their surface binds to the Angiotensin-converting enzyme 2 (ACE-2) receptor found in abundance in the olfactory region (OR) (Chen, Shen et al. 2020, Chavda, Vora et al. 2021, Pilicheva and Boyuklieva 2021). This might be the reason for loss of olfaction among COVID-19 patients and suggests intranasal therapy.

Intranasal vaccines seem promising because they function similarly to viral infection. Additionally, they can be conveniently and painlessly self-administered (Chavda, Vora et al. 2021). From the immunology perspective, intranasal vaccines trigger both mucosal and systemic immune responses (Krammer 2020, Chavda, Vora et al. 2021). Experimental studies demonstrate that neutralizing SARS-CoV-2-specific mucosal IgA antibodies could vanish the infection (Butler, Crowley et al. 2021, Chavda, Vora et al. 2021). However, intramuscular vaccines usually can only trigger robust IgG response in the lower respiratory tract, but cannot lead to adequate mucosal IgA response in the upper respiratory tract. Therefore, systemically vaccinated individuals are still susceptible to asymptomatic viral infection and can transmit the live virus to others (Bleier, Ramanathan et al. 2020). Hence, nasal spray delivery of SARS-CoV-2 vaccine candidates to ACE-2-rich areas can be substantially effective in preventing COVID-19 infection and transmission (Xi, Lei et al. 2021).

Other than the benefits mentioned above for intranasal vaccines, they can also benefit individuals and children who are afraid of needles (Chavda, Vora et al. 2021). It was reported (Flaherty 2021) that children and adolescents could be dangerous carriers and spreaders of COVID-19 due to their capability of carrying high levels of SARS-CoV-2 in their respiratory secretions. Preventing SARS-CoV-2 transmission and infection among them will be highly beneficial for community immunity (Glezen 2001). Therefore, it is necessary to investigate how intranasal vaccine transport and triggers the immune system response in the pulmonary routes of children.

To administer intranasal vaccines with desired effectiveness, some factors need to be fine-tuned. Among all the factors mentioned in the literature (Xi, Lei et al. 2021), nasal ACE-2-rich locations as the targeted delivery sites, the type of nasal spray device, delivery strategy, nasal physiology, and inter-subject variability are possibly the main factors in determining the vaccination effectiveness. Specifically, ACE-2-rich areas in the nasal cavity should be the target site for vaccine delivery. However, the complex morphology of the nasal passage makes it challenging for the vaccine droplets to reach the targeted area. In addition, inter-subject variability of the nasal cavity anatomy between different age groups (e.g., adults vs. children) can significantly influence the transport and deposition of inhaled nasal spray droplets. Therefore, assessment of nasal vaccine delivery for children using their age-specific nasal cavity geometries is necessary since it will be able to avoid biased conclusions drawn from the research results obtained *via* using adult nasal cavity geometries.

As of December 2021, There has been some nasal vaccine sprays Phase 1 clinical trial against SARS-CoV-2 (Hassan, Kafai et al. 2020). A fully preventative spray (Moakes, Davies et al. 2021) was formulated to target the lining of the upper respiratory system against SARS-CoV-2 as an intranasal vaccine potential candidate. It is a polysaccharide-based spray, of which the physical properties, spray droplet size distribution, and antiviral properties were documented. In detail, gellan gum (GG) and λ-carrageenan (λ-C) are two biocompatible and intrinsically mucoadhesive components of vaccine solution. Their high viscosity can reduce the clearance due to dripping, thereby enhancing spray residence time in the nasal cavity (Moakes, Davies et al. 2021). It has also been claimed that GG and λ-C demonstrate antiviral capacities. Phosphate-buffered saline (PBS) was added to the vaccine solution because of its buffering capacity against GG and λ-C native acid pH. It is possible to deliver the preventative spray (Moakes, Davies et al. 2021) as the intranasal vaccine to the OR to cover more epithelial cells and minimize the available receptors for SARS-CoV-2.

There have been many CFPD studies elucidating the transport dynamics of spray particles/droplets in the nasal cavity in the past two decades, focusing on the targeted drug delivery to the OR and other regions in the main nasal airway passage. Inthavong, Tian et al. (2007), Inthavong, Tian et al. (2008), and Tong, Dong et al. (2016) investigated the delivery of nasal spray particles (with selective sizes) into the nasal cavity of a 25-year-old male. The above-mentioned papers investigated particle deposition in the main passage of the nasal cavity, but did not target OR. Kiaee, Wachtel et al. (2018) studied nasal spray particle transport in 7 realistic adult nasal airway geometries. Based on their outcomes, nasal spray particle penetration through the nasal cavity is highly sensitive to particle size. They found particles with 20 μm – 30 μm in diameter are preferred since their deposition in OR is the highest. Calmet, Inthavong et al. (2019) simulated nasal spray particle deposition in a human nasal cavity under multiple inhalation conditions. They claimed that spray deposition in OR is negligible and unaffected by the spray particle conditions, i.e., spray cone angle, insertion angle, and initial velocity. Zare, Aalaei et al. (2022) studied drug delivery to the inferior meatus of the nasal airway model of a 42-year-old female using a spray device with an angled tip. Drug particles were found to be guided toward the target area more efficiently. None of the computational efforts mentioned above has focused on aerosolized COVID-19 vaccine candidates or systematically investigated how administration strategies can influence the deposition of vaccine droplets in the nasal cavity of a young child with realistic relative humidity (RH) and temperature distribution. Additionally, although there are research efforts on modeling SARS-CoV-2 induced immune system responses (Li, Kuga et al. 2021, Vaidya, Bloomquist et al. 2021), dynamics of both innate and adaptive immune system responses against SARS-CoV-2 have not been modeled.

Therefore, focusing on children, this study predicted transport, deposition, and the immune system response of COVID-19 nasal vaccine spray droplets in an upper airway model of a six-year-old child covering from the nasal cavity to generation 5 (G5) using a multiscale numerical model, i.e., CFPD-HCD model. The CFPD-HCD model is experimentally calibrated and validated. The transport and deposition of polydisperse COVID-19 nasal vaccine multicomponent droplets with initial diameters ranging from 20 μm to 300 μm were simulated using an experimentally validated CFPD model, which also considered the evaporation/condensation effect on the droplet size change. The vaccine solution consists of water, PBS, GG, and λ-C (Moakes, Davies et al. 2021). Parameters that can influence the delivery efficiency of the intranasal COVID-19 vaccine to the targeted ACE-2-rich region, i.e., the olfactory region (OR), were determined based on the numerical simulation results reported in the literature (Kiaee, Wachtel et al. 2018, Xi, Lei et al. 2021). Those parameters include spray cone angle, spray droplet initial velocity, and initial droplet composition. Furthermore, the vaccine droplet deposition data were converted as the initial conditions for the HCD model to predict the variations in immune system responses based on different intranasal administration strategies for the COVID-19 nasal spray vaccine. For this purpose, vaccine coverage (VC) was calculated, which is defined as the percentage of epithelial cells in OR which can be covered by vaccine droplets. Based on the VC value, epithelial cell counts that are susceptible to infection were obtained and introduced to the HCD model. In the present HCD model, the kinetics of macrophages and natural killer (NK) kinetics were appended to the model introduced by (Lee, Topham et al. 2009).

## 5 Governing equations

### 2.1 Computational fluid-particle dynamics (CFPD) model

An experimentally validated CFPD model based on the one-way coupled Euler-Lagrange method was used to simulate multicomponent droplet transport in the respiratory system model (Hayati, Feng et al. 2021). It was assumed that the droplets are spherical, and droplet-droplet interaction was neglected. Droplets are composed of water, GG, λ-G, and PBS (Moakes, Davies et al. 2021). Water is the only evaporable component. Nasal mucosa was not explicitly modeled as a separate fluid phase in this study. Instead, the 100% trapped wall boundary condition was applied, assuming droplets will deposit when touching the airway walls.

#### 2.1.1 Humid airflow

Since the airflow regime in the respiratory system model was transitional between laminar and turbulence, the transition shear stress transport (SST) (Menter, Langtry et al. 2006) model was employed to capture the laminar-to-turbulent transition sites accurately. The conservation laws of mass, momentum, energy, turbulence kinetic energy (*k*), and the specific rate of dissipation (*ω*) are provided in previous studies (Haghnegahdar, Zhao et al., 2019). The convection-diffusion equation (Bird, Stewart et al. 1960) is employed for the calculation of water vapor (*y_w_*) distribution to predict droplet size change dynamics, i.e.,

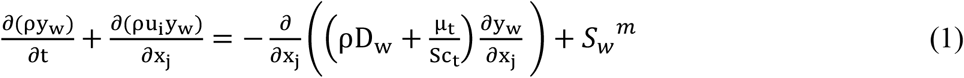

In this study, air and water vapor are the constituents of the gas mixture. Accordingly, ρ is the density of the mixture, Dw is the molecular diffusivity of water in the air, μt is the turbulent viscosity, and *S_w_^m^* is the mass source term (i.e., the evaporation rate of water between humid air and droplet).

The energy equation (Longest and Xi 2008) was solved to predict the temperature distribution in the computational domain, i.e.,

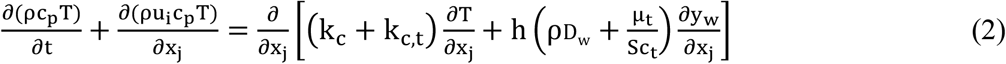

where c_p_, T, k_c_, and k_c,t_ are the specific heat of the gas mixture, temperature, thermal conductivity, and turbulent thermal conductivity, respectively. In addition, h is the convective heat transfer coefficient of water. Please refer to Hayati, Feng et al. (2021) for more details on the governing equations.

#### 2.1.2 Multicomponent droplet transport

Droplets traveling along with fluid flow are subjected to multiple forces. To obtain the droplet trajectories, Newton’s Second Law was solved (Hayati, Soltani Goharrizi et al. 2019, Hayati, Feng et al. 2021), i.e.,

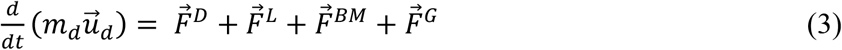

where 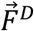 is the drag force (Chen, Feng et al. 2017), 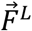 is the Saffman lift force (Saffman 1965), 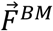 is the Brownian motion induced force (Hayati, Soltani Goharrizi et al. 2019), and 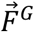 is gravity.

Droplet mass and energy balance equations is employed to calculate the size change induced by evaporation and condensation. The mass balance equation for droplets can be given as

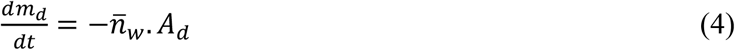

where 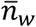 is the average mass flux of water evaporation/condensation on the droplet surface, which can be defined by

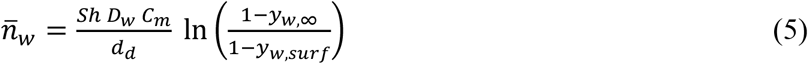

In Eq. (5), *y*_*w*,∞_ and *y_w,surf_* are the mass fractions of water in the vapor phase and at the surface of droplet, respectively. D_w_ is the mass diffusivity of water, *Sh* is the Sherwood number (Whitaker 1972), and C_m_ is the Fuchs-Knudsen correction factor (Chen, Feng et al. 2017).

The Sherwood number (*Sh*) can be calculated by

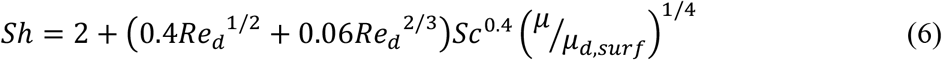

where Sc=μ/ρD_e_ is the Schmidt number, Re_d_ is the droplet Reynolds number, μ and μ_d,surf_ are gas mixture viscosities in the fluid and on the droplet surface. This study assumes *^μ^/μ_d,surf_*=1 (Whitaker 1972).

Furthermore, C_m_ can be given as

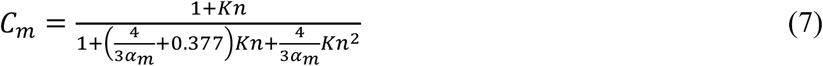

where Kn is the Knudsen number, and α_m_=1 is the mass accommodation factor (Broday and Georgopoulos 2001).

It is worth mentioning that the Kelvin effect was taken into account when calculating the water mass fraction *y_w,surf_* at the saturated droplet surface in Eq. (5). Specifically, compared with flat liquid surface, the droplet surface curvature allows water molecules to evaporate more freely and faster. Such a phenomenon, i.e., the Kelvin effect, causes the equilibrium vapor pressure to be higher than the saturation vapor pressure at the droplet surface (Brechtel and Kreidenweis 1999, Asgharian 2004). Therefore, the Kelvin effect factor *K_w_* can be defined and calculated by

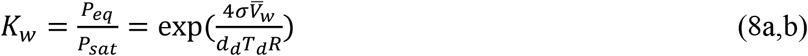

Here, P_sat_ is saturation vapor pressure at droplet temperature and P_eq_ is equilibrium vapor pressure above the droplet. σ denotes droplet surface tension, R=8.314 J/mol-k is the gas constant, and 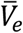 is the molar volume of evaporable component, which is defined by (Brechtel and Kreidenweis 1999)

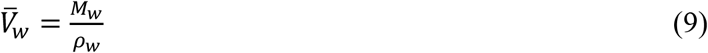

where M_w_ and ρ_w_ are molecular weight and density (Kreidenweis, Koehler et al. 2005) of water, respectively. Therefore, *y_w,surf_* should be calculated by

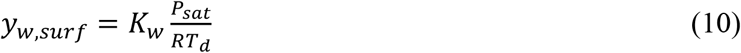

The energy balance equation for droplets can be given as

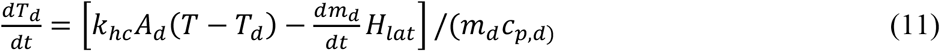

where *H_lat_* is the latent heat. T_d_ and c_p,d_ are droplet temperature and specific heat. *k_hc_* denotes modified thermal conductivity, which can be calculated by

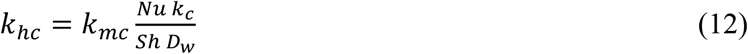

In Eq. (12), k_c_ is the thermal activity of humid air, Nu is the Nusselt number, and k_mc_=C_m_D_w_Sh/d_d_ is the mass transfer coefficient. Assuming there is no internal resistance to heat transfer inside the droplet, Nu can be defined by (Whitaker 1972)

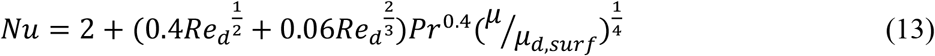

### 2.2 Host cell dynamics (HCD) model

In this study, a two-compartment modeling framework system (i.e., a lung compartment and a lymphatic compartment) of 24 coupled nonlinear stiff ordinary differential equations (ODEs) introduced by Lee, Topham et al. (2009) is developed and revised to model the immune system response. The HCD model includes parameters representing the rates at which the viral titer, epithelial cells, and immune cell counts vary. Specifically, viral titer, epithelial cells, interferons, macrophages, NK cells, and dendritic cells are modeled and studied under the lung compartment. Virus-loaded dendritic cells, T cells, B cells, effector cells, and antibodies are simulated and analyzed as the lymphatic compartment. The role of the adaptive effector cells and antibodies are considered in mitigating infected cells and viral load in the lung compartment, respectively. The ODEs of the HCD model and more details can be found in Appendix. Compared with existing HCD models for SARS-CoV-2 study (Li, Kuga et al. 2021, Vaidya, Bloomquist et al. 2021), the HCD model developed and employed in this study is more advanced since it includes the kinetics of both innate (macrophages, NK cells, interferons, and dendritic cells) and adaptive (CD8^+^ T cells, short-lived, and long-lived antibodies) immune system responses.

## 6 Initial and boundary conditions

To use the realistic temperature and RH initial conditions inside the respiratory system model before intranasal vaccine spray administration, one breathing cycle was simulated to obtain the temperature and water vapor mass fraction distributions. Details of the boundary conditions are listed in Table 1. Temperature and humidity distribution at airway walls were adopted from Ferron, Upadhyay et al. (2013) for the one-breathing cycle simulations before vaccine administration. Vaccine droplets were sprayed into the nasal cavity once the exhalation phase ended.

**Table 1.**
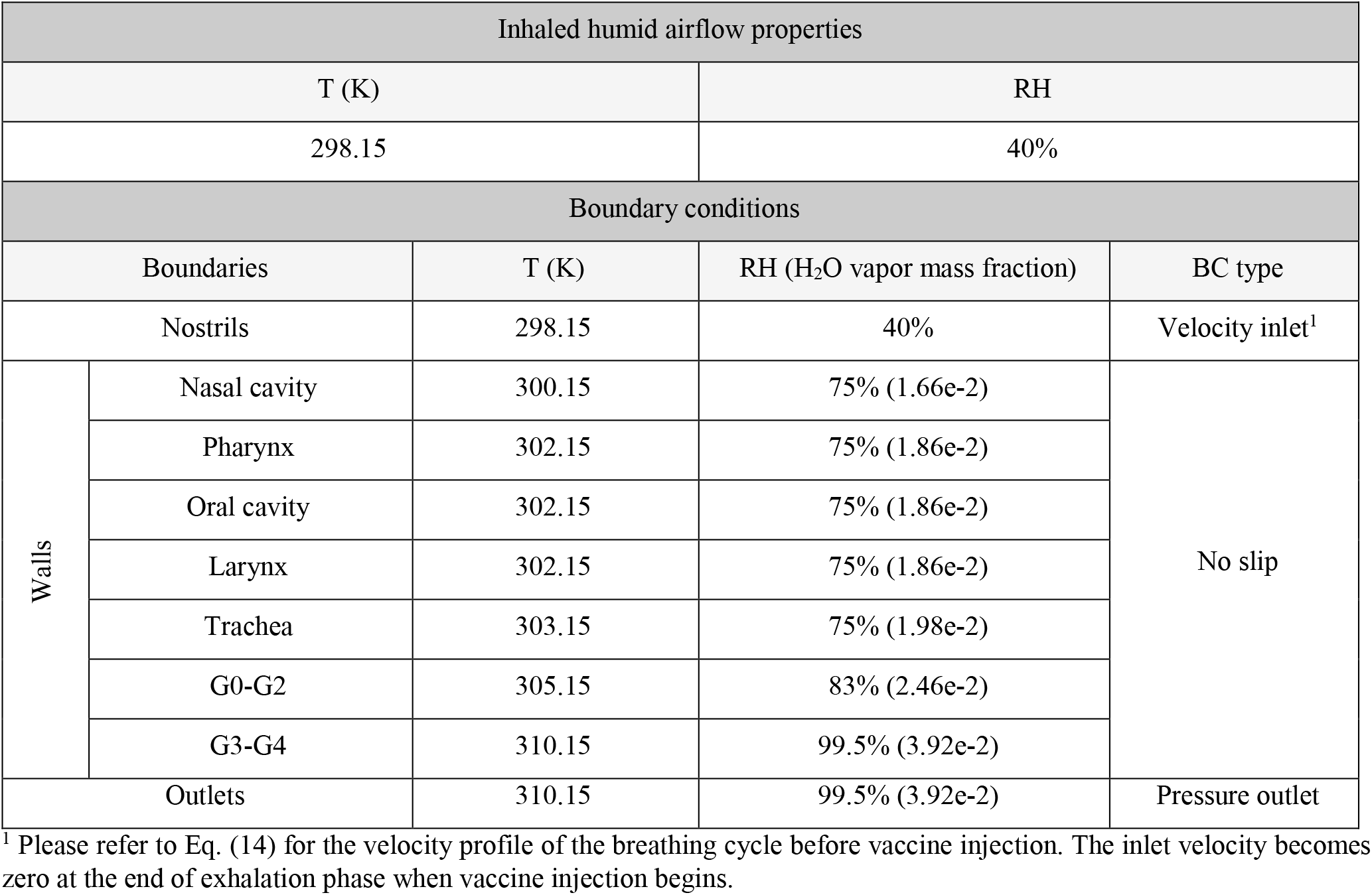
Airflow properties and boundary conditions

Since patients usually do not breathe when spraying, the airflow inlet velocity is assumed to be zero when injecting vaccine droplets. Droplet properties and compositions (Moakes, Davies et al. 2021) are presented in Table 2.

**Table 2.** Droplet compositions and properties

| Droplets properties |  |  |  |  |  |  |  |  |
| --- | --- | --- | --- | --- | --- | --- | --- | --- |
| Components | Components mass fraction |  |  |  |  |  |  |  |
| | GG to λ-C mass ratio = 75:25 | | GG to λ-C mass ratio = 50:50 | | GG to λ-C mass ratio =25:75 | | | |
|  | Case 1<br>0.2%<sup>1</sup> | Case 2<br>1% | Case 3<br>1% |  | Case 4<br>1% |  |  |  |
| PBS | 4.70e-2 | 4.66e-2 | 4.66e-2 |  | 4.66e-2 |  |  |  |
| GG | 1.43e-3 | 7.08e-3 | 4.72e-3 |  | 2.36e-3 |  |  |  |
| λ-C | 4.75e-4 | 2.36e-3 | 4.72e-3 | | 7.08e-3 | | | |
| Water | 9.51e-1 | 9.44e-1 | 9.44e-1 |  | 9.44e-1 |  |  |  |
| T (k) | MW (g/mol) |  |  |  | Density (g/cm<sup>3</sup>) |  |  |  |
| | PBS | GG | λ-C | water | PBS | GG<sup>2</sup> | λ-C<sup>3</sup> | Water |
| 298.15 | 65.22 | 128.61 | 579.5 | 18.01 | 2.09 | 0.5 | 0.034 | 1.00 |

<sup>1</sup>Total polymer concentration (%w/v)
<sup>2</sup>Morrison, Talashek et al. (2016)
<sup>3</sup>Density of vaccine solution was reported 1010 kg/m<sup>3</sup> by Moakes, Davies et al. (2021). Density of λ-C was calculated accordingly.

Initial conditions for HCD model calibration are mentioned in Table A3 in the Appendix. There are approximately 4e8 epithelial cells in the upper respiratory tract of humans (Baccam, Beauchemin et al. 2006). It was also estimated that 1% to 20% of these cells express ACE2 receptors (Hou, Okuda et al. 2020, Sungnak, Huang et al. 2020). In this study, it was assumed that 10% of the 4e8 epithelial cells have ACE-2 and are susceptible to being infected by SARS-CoV-2 (Vaidya, Bloomquist et al. 2021). Since there was no available clinical data for young children, the initial number of susceptible epithelial cells with ACE-2 to SARS-CoV-2 infection is E_0_=4e7 that was reported for adults was employed for the 6-year-old child. Based on the clinical samples, the viral titer was VT_0_=2e5 copies/ml on day 0 after symptoms onset (Pan, Chen et al. 2020). Therefore, VT_0_=2e5 copies/ml was also used for the HCD simulations in this study. Initial values counts for immature dendritic cells, naive CD4+ T cells, and naive B cells were considered 1000 cells (Lee, Topham et al. 2009). The initial counts of uninfected macrophages and NK cells residing in the lung compartment are assumed to be 1000 cells as well. The initial counts of other cell counts were set to zeros. After HCD model calibration, E0 was updated based on the vaccine coverage of ACE2 in OR obtained by CFPD analysis.

## 7 Numerical Simulation

### 4.1 Geometry and mesh

The respiratory system model employed for the current study is shown in Fig. 1(a). The model comprises the entire upper airways (nasal cavity, oral cavity, pharynx, larynx, and trachea) and the first five generations (G) (i.e., G0-G4) of the tracheobronchial (TB) tree. Specifically, the larynx-to-trachea region was reconstructed from the CT scan images of a 6-year-old female. The mouth, nose, nasopharynx, and pharynx were adopted from a subject-specific upper airway geometry reconstructed from the magnetic resonance imaging (MRI) data of a 47-year-old healthy male. The geometry were scaled down in size to be compatible with the anatomical geometry dimensions for the 6-year-old age group. The G0-G4 TB tree starting from G0 to G4 was generated with anatomical features using a stochastic algorithm (Kitaoka 2011). The surface areas of different regions of the geometry are listed Table 3.

**Fig. 1.**
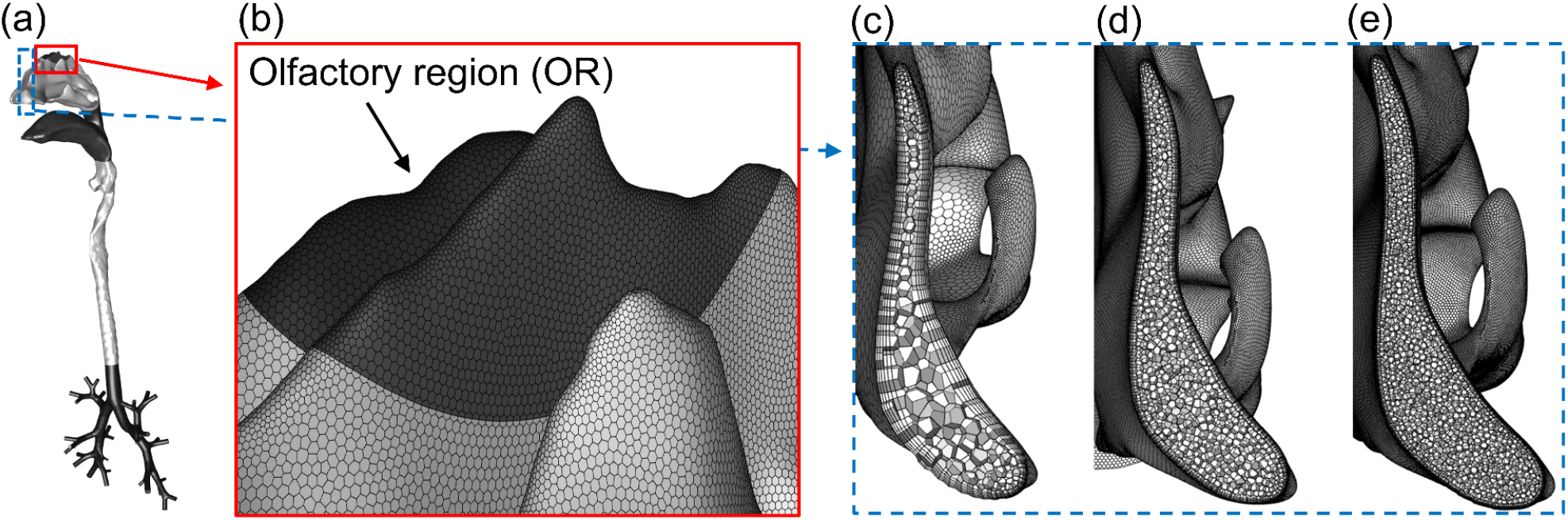
Respiratory system geometry and mesh: (a) the upper airway geometry from nasal cavity to G4 of a 6-year old child; (b) the surface mesh of the olfactory region (i.e., the dark gray region). Volume mesh details of (c) Mesh 1 (coarse); (d) Mesh 2 (final mesh); and (e) Mesh 3 (most refined).

**Table 3.** Surface area of different regions in the respiratory system model

|  | Respiratory system model | Nasal cavity | Olfactory regions |
| --- | --- | --- | --- |
| Surface area<br>(m<sup>2</sup>) | 13.1e-3 | 47.05e-4 | 17.8e-5 |

Unstructured polyhedron-based meshes were generated for the upper airway geometry using Ansys Fluent Meshing 2020 R1 (Ansys Inc., Canonsburg, PA). Five near-wall prism layers were generated to precisely capture boundary layer flow regimes and laminar-to-turbulence transition sites. Details of the surface mesh are shown in Fig. 1 (b) as well as the OR (in dark gray). To find the final mesh that can provide the optimal balance between computational efficiency and accuracy, three meshes with different element sizes were generated for the mesh independence test. Mesh specifications are listed in Table 4. Polyhedron-base volume cells of the meshes in a cross-section of the nasal cavity are visualized in Figs. 1 (c)-(e).

**Table 4.** Specifications of meshes generated for the independence test

| Mesh number | Cell counts | Iterations needed for convergence | $y^+$ |
| --- | --- | --- | --- |
| Mesh 1 | 1,150,080 | 131 | 98 % < 1 |
| Mesh 2 (Final Mesh) | 3,725,651 | 241 | 99 % < 1 |
| Mesh 3 | 6,550,677 | 375 | 100 % < 1 |

The normal breathing frequency of a 6-year-old child was considered 25 breaths/min (Fleming,Thompson et al. 2011), and the tidal volume for a 21 kg child (average weight of children at 6 years old) was considered 7 ml/kg (Koomen, Nijman et al. 2021). Employing the above-mentioned specifications, the steady-state inlet flow velocity employed at each nostril for the mesh independence test was 4.5 m/s (i.e., the average inhalation velocity). Using the average inhalation velocity, an idealized sinusoidal breathing waveform was employed for this study. The transient velocity V at the nostrils is defined as a function of time t, i.e.,

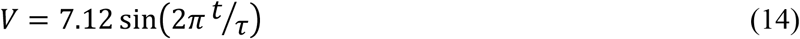

where τ=2.4 s is the duration of one breathing cycle.

The convergence criteria were set to 1e-4. The SIMPLE scheme was employed for pressure-velocity coupling, and the first-order upwind scheme was used for the spatial discretization of pressure, turbulent kinetic energy, and specific dissipation rate. Figures 2 (a)-(c) show the comparisons of the velocity profiles along three lines selected at different locations in the respiratory system. The comparison indicates that Mesh 2 is the most efficient mesh to obtain numerical results with acceptable accuracy.

**Fig. 2.**
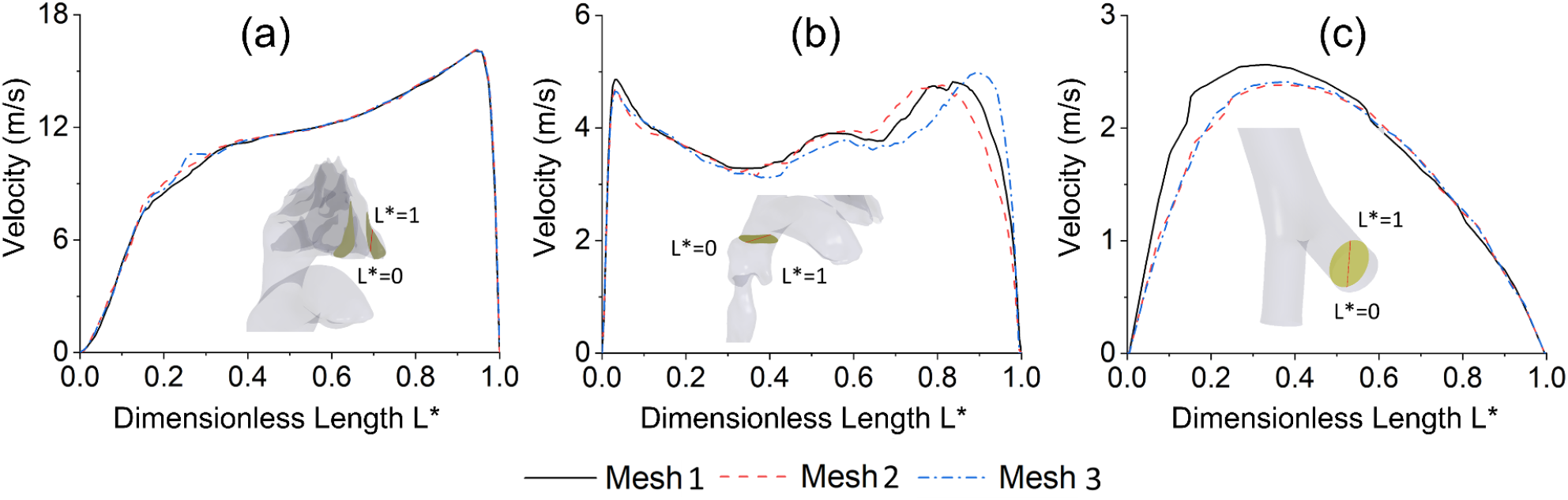
Airflow velocity profile comparisons using multiple meshes along with selected lines in: (a) nasal cavity, (b) larynx, and (c) G4.

### 4.2 CFPD setup

Ansys Fluent 2022 R1 (Ansys Inc., Canonsburg, PA) was utilized for CFPD simulations. The flow field is transient with the flow time step size Δ*t_f_* = 2e-3 s, which is determined by a time-step independence test. The transient breathing waveform (i.e., Eq. (14)) was applied at the nostrils. As mentioned in Section 3, vaccine droplets were injected into the left nasal cavity after one breathing cycle simulated, with the initial velocity equal to 8.5±3.5 m/s (Xi, Lei et al. 2021). Droplet size distribution was considered polydisperse ranging from 20 μm to 300 μm (Xi, Lei et al. 2021), and 40 different size bins (40,000 particles) were set to be injected. The spray nozzle diameter was 0.2 mm (Kapadia, Grullo et al. 2019), and the insertion depth was 5 mm from the nostril. The spray nozzle was set to make an angle of 35° with the gravitational direction, assuming that the head-to-foot direction is aligned with gravity.

The droplet time step size is 1e-4 s. Constant binary diffusivity (i.e., *D_w_*=5.05e-5 m^2^/s) and piecewise-linear saturation vapor pressure were set for the liquid water component in vaccine droplets. Therefore, the saturation vapor pressure is temperature-dependent and is defined as

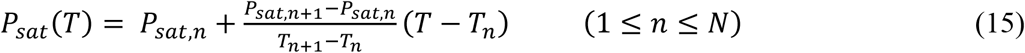

where *N* is the number of segments between maximum and minimum temperature. Saturation vapor pressure data used in Eq. (15) were obtained from an engineering handbook (Perry, Gree et al. 1984).

Furthermore, in-house user-defined functions (UDFs) were revised and compiled in the computational cases for

a. Transient inhalation and exhalation profile;
b. Droplet Brownian motion induced force;
c. Recovering anisotropic near-wall turbulence fluctuation velocities and drag coefficient;
d. Calculating multicomponent droplet size change;
e. Updating droplet density;
f. Storing locations and physical properties of droplets deposited; and
g. Storing the surface area of the faces droplets landed on.

Numerical simulations were performed on a local Dell Precision T7810 workstation (Intel^®^ Xeon^®^ Processor E5-2643 v4 with dual processors, 64 cores, and 128 GB RAM) and a local Dell Precision T7910 workstation (Intel^®^Xeon^®^ Processor E5-2683 v4 with dual processors, 64 cores, and 256 GB RAM). Using 32 cores, the CPU time for the simulation of one breathing cycle to initialize the case was approximately 75 hours, and the nasal spray droplet transport simulations took approximately 0.05 hours.

### 4.3 HCD setup

To solve the system of stiff nonlinear coupled ODEs that can quantitatively describe immune response dynamics, an in-house MATLAB code was developed. An ODE solver (i.e., ode23t), capable of using adaptive time step size for resolving nonlinear stiff ODEs, was employed. The initial values of the variables are provided in Table A3 in the Appendix. Temporal behavior and dynamics of the viral titer and innate immunity were modeled in the lung compartment. Dendritic cell maturation, naïve T cell activation, and antibodies and effector cell proliferation are simulated in the lymphatic compartment. Two and a half (i.e., 2.5) days after the onset of the symptoms, the antigen-specific antibodies are detectable in the lung compartment (Long, Liu et al. 2020). Therefore, a delay time, *τ_D_*=1 day, was considered to take into account the time needed for adaptive immune response to SARS-CoV-2 to start being activated, and another delay time, *τ_T_*=2 days, was included in the optimization to observe the adaptive immunity dynamics in the lung compartment. The simulation was performed for 30 days in the lung compartment and 28 days (from day 1 to day 29) in the lymphatic compartment. The HCD model coefficients were well-calibrated by minimizing the root mean squared error (RMSE) between available clinical data and HCD modeling results.

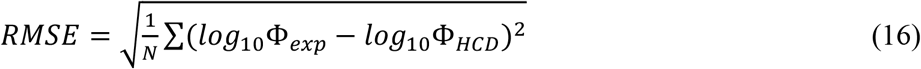

where, Φ_exp_ and Φ_HCD_ are experimental and computational data, respectively.

The details of the HCD model calibrations are discussed in Section 5.2. After calibration, the initial epithelial cell counts E0 obtained from CFPD simulation results were introduced to the HCD model for immune response analysis after intranasal vaccine injection (please see Section 3 for more details).

## 8 Model validation and calibration

### 5.1 CFPD model validation

To validate the CFPD model, pure water droplet evaporation was simulated and compared with experimental data. Specifically, a 16-μm pure water droplet was released into a square duct. The geometric dimensions of the duct are 0.15 m in height × 0.15 m in width and 5.5 m in length, respectively. Ambient relative humidity (RH) and temperature were 70% and 296.15 K constantly in the duct. Inlet air velocity was 1 m/s. Droplet initial temperature was set to 296.15 K, and its initial velocity was 0 m/s. Figure 3 compares the simulation results and experimental data reported by El Golli, Bricard et al. (1974) on the droplet radius change. Good agreement can be observed between numerical simulation and experimental measurements, indicating that the CFPD model employed in this study can accurately predict the water evaporation/condensation rate in the vaccine droplets simulations.

**Fig. 3.**
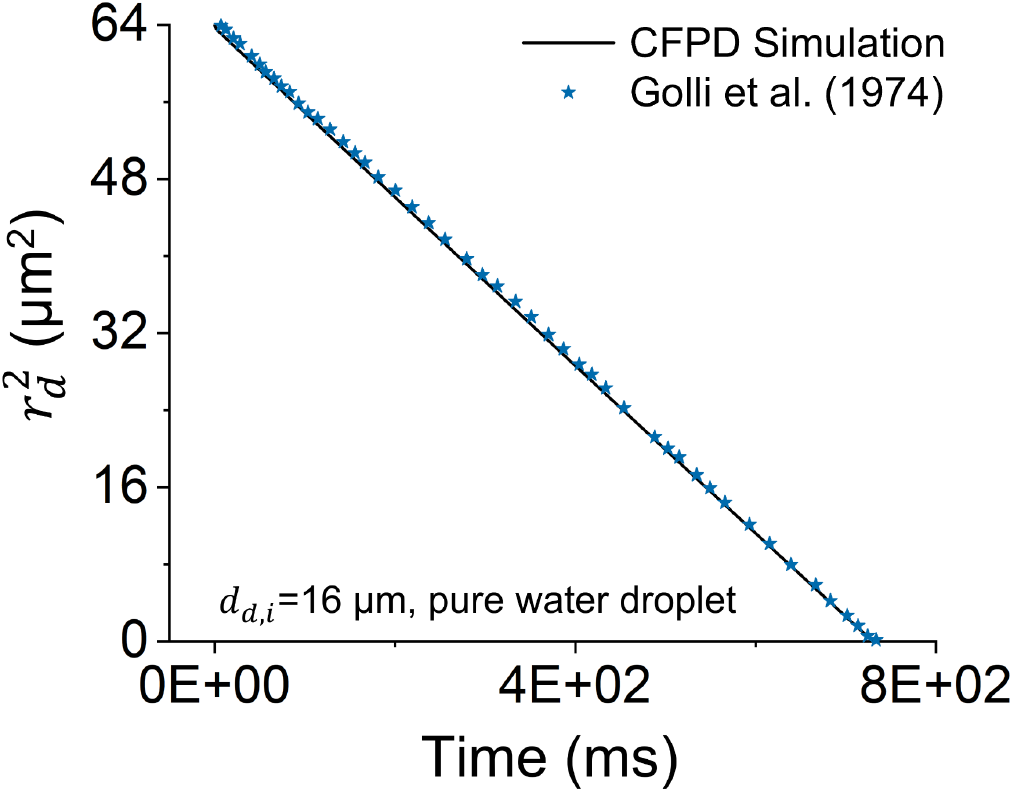
CFPD model validation for pure water droplet evaporation

### 5.2 HCD model calibration

Calibrated with clinical data obtained from human patients who were tested positive for SARS-CoV-2, the code estimated the optimized values for the viral load and the immune response coefficients. For optimizing the coefficients, the “Multiobjective optimization using Genetic Algorithm” (i.e., gamultiobj in MATLAB) was employed. Specifically, a set of coefficient values were computed using the above-mentioned optimization algorithm by minimizing the RMSE magnitude between the experimental data sets obtained from COVID-19 patients (Long, Liu et al. 2020, Pan, Chen et al. 2020) and the corresponding data predicted by the HCD model (see Eq. (16)). Through the optimization, some coefficient values were considered fixed (see Appendix for more details). The coefficient values optimized are listed in Table A4 in the Appendix.

Comparisons of clinical data and the HCD simulation results after calibration are shown in Fig. 4. Specifically, the fitted curves for viral load and adaptive immune responses for a baseline simulation were compared. RMSE values for cell count curves predicted by the HCD model are reported as well. It can be observed that the HCD model predictions are in good agreement with clinical data on viral clearance and antigen-specific antibody titer time profile. Viral load peaks approximately 2 to 3 days after symptoms onset (Fig. 4 (a)), which suggests virus production rate per infected epithelial cell is *π_V_* = 2.61 copies/(ml day). Furthermore, the death rate constant for infected cells is *δ_EI_*=5 /day. Viral load starts decreasing at a high rate after peak day mainly because the number of healthy epithelial cells drops, and infectious viruses do not have an environment to replicate themselves in, i.e., virus non-specific clearance. It is shown in Figs. 4 (b) and (c) that SARS-CoV-2 specific antibodies (IgM and IgG) are not detectable in the lung compartment before Day 2 after symptoms onset. After two days, however, the antibodies IgM and IgG travel to the lung from the lymph node to neutralize the viruses at the rate of *c_AS_*= 0.53 /(day μg) and *c_AL_*=1.1 /(day μg). Both IgM and IgG levels plateaued after day 15.

**Fig. 4.**
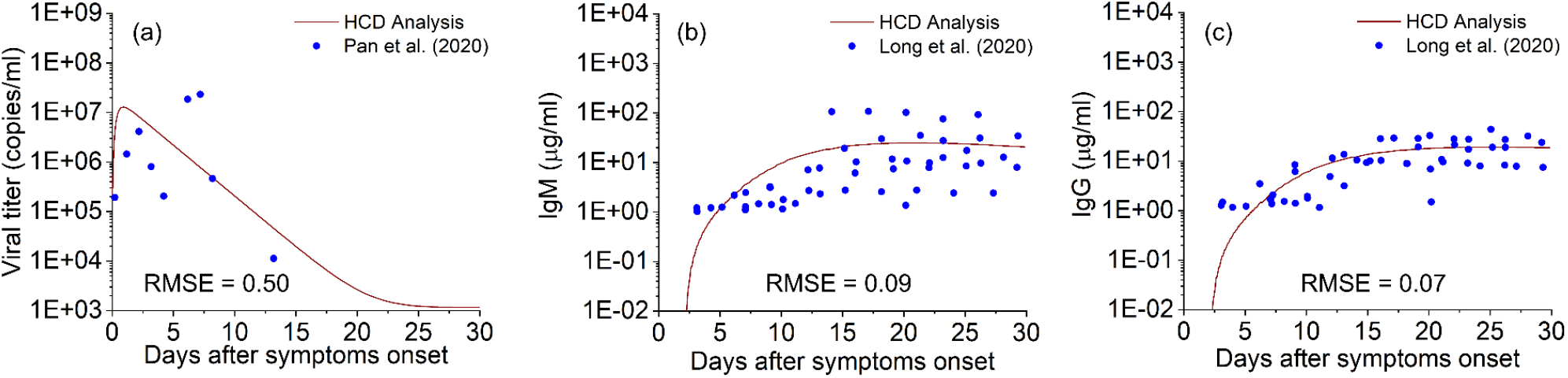
Comparisons of optimized HCD modeling results and available clinical data to predict SARS-CoV-2 infection induced temporal variations in (a) viral titer, (b) short-lived antibodies (IgM), and (c) long-lived antibodies (IgG)

## 9 Results and discussion

### 6.1 RH distribution in the nasal cavity

Figure 5 shows the water vapor distribution in the nasal cavity at the end of the exhalation phase when the airflow velocity is zero and vaccine spraying begins. Local values of minimum and maximum water vapor mass fraction are reported under and above each cross-section. Besides the closest cross-section to the nostril, the difference between the maximum and minimum water vapor mass fraction at each cross-section is small, indicating the effectiveness of humidification of the nasal passage to the inhaled dry air.

**Fig. 5.**
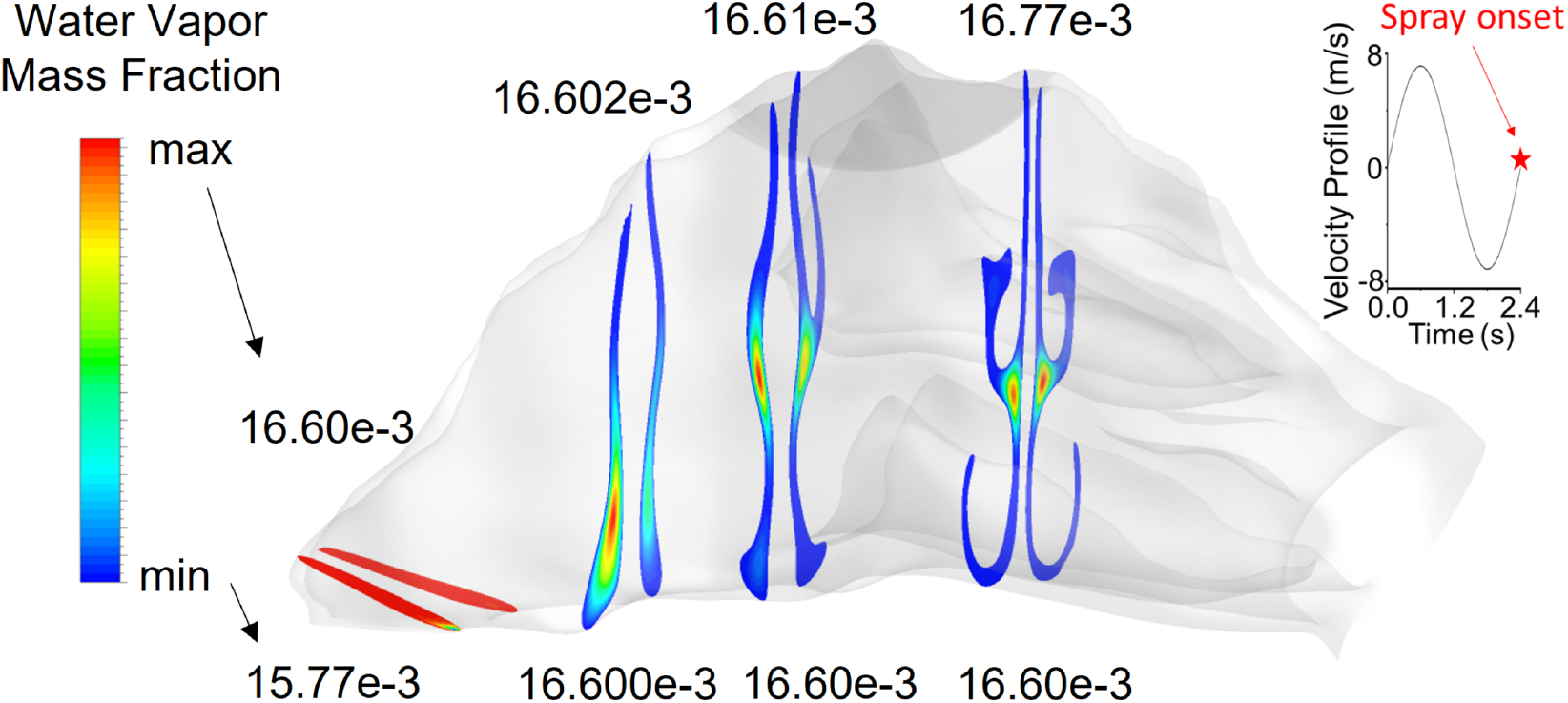
Mass fraction distribution of water vapor in multiple cross-sections of the nasal cavity at the end of exhalation when vaccine droplets were injected.

### 6.2 Influence of multiple parameters on vaccine delivery efficiency to the olfactory region

To find the key parameters that can influence the delivery efficiency of vaccine droplets to the OR, numerical simulations were performed with various initial spray droplet velocities (*V_d,i_*), spray cone angles (θ), and initial droplet compositions. Figure 6 demonstrates the delivery efficiency of vaccine droplets to OR, influenced by the above-mentioned parameters.

**Fig. 6.**
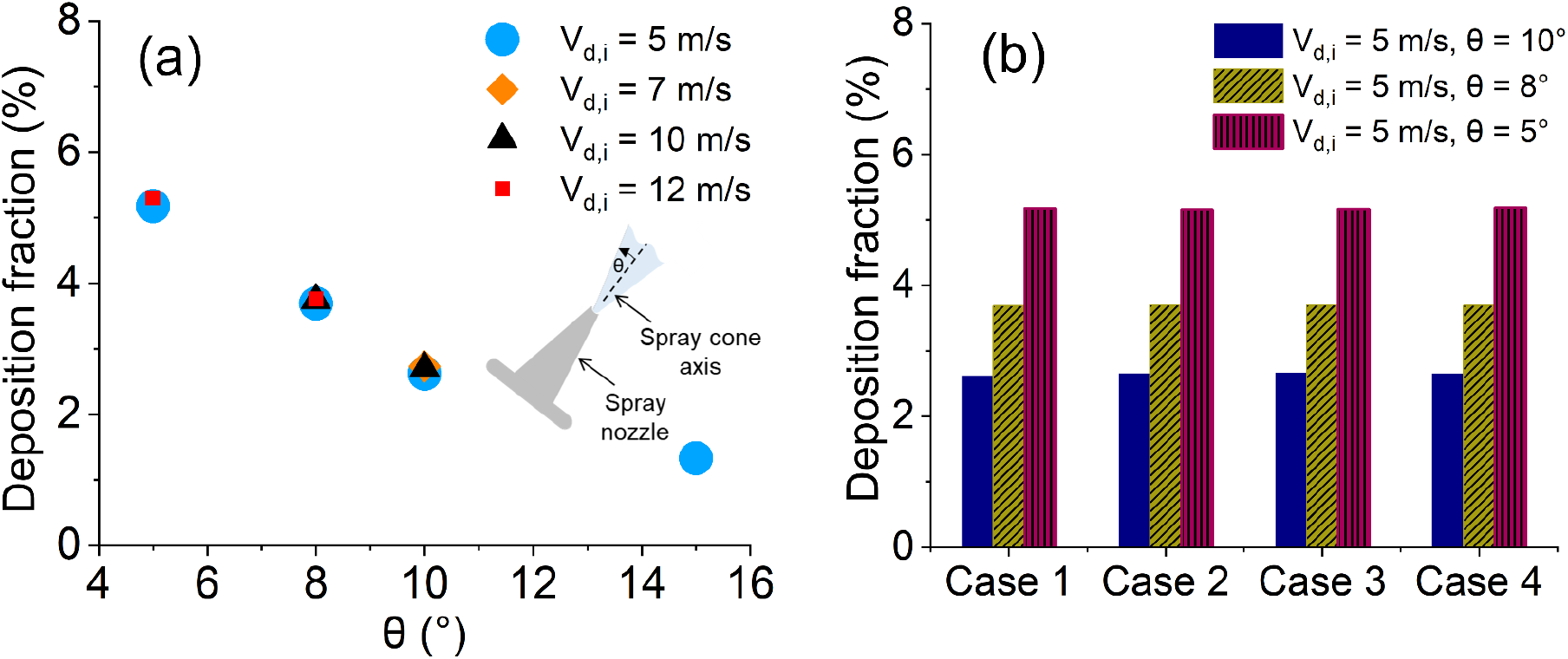
Deposition fractions (DF) of the nasal spray vaccine droplets in the olfactory region (OR)

Specifically, Fig. 6 (a) shows the deposition fraction (DF) of droplets versus spray cone angle θ for four different droplet initial velocities *V_d,i_*. It can be observed that with the same *V_d,i_*, droplet DF in OR decreases with the increase in θ. For example, with *V_d,i_* = 5 m/s, DF for θ = 5° is approximately 5%, while DF decreases lower than 1.5% for θ = 15°. Therefore, Fig. 6 (a) indicates that a smaller spray angle is recommended for higher vaccine delivery efficiency to the OR for the 6-year-old child. The reason for this trend is that with a wider spray cone angle, more micro-sized droplets can intercept the airway wall due to the complex morphology and narrow nasal airway passage. Furthermore, Fig. 6 (a) also shows that varying *V_d,i_* has a negligible impact on the vaccine droplet DF in OR, which is because the initial droplet velocity (*V_d,i_*= 5 m/s to 12 m/s) and momentum are relatively high, the viscous dissipation effect induced by the drag force is insignificant. In addition, due to the small droplet diameter, the gravitational sedimentation effect on droplet trajectory is negligible compared with the high initial momentum of the droplets. Therefore, although the droplet residence time was influenced by *Vd,i*, the droplet trajectories and deposition locations are highly similar for the droplets released at the same position with the same θ and different *V_d,i_*.

Figure 6 (b) shows regional DFs of vaccine droplets in OR with different droplet initial compositions (see Table 3 for the initial droplet composition associated with each case) and injection cone angles θ (5°, 8°, and 10°). Since *V_d,i_* has negligible influence on droplet DF in OR, *V_d,i_* was kept constant at 5 m/s in Fig. 6 (b). Figure 6 (b) demonstrates again that the wider cone angle leads to lower vaccine delivery efficiency to OR. In contrast, droplet composition does not significantly influence the DF of vaccine droplets to the targeted site (i.e., OR). For all four compositions, water liquid mass fraction is approximately 95% in the droplet, while other non-evaporable components constitute only 5% in mass. Although the densities of the non-evaporable components are different, the variation in their mass fraction among Cases 1 to 4 (see Table 2) is small. Hence, there will not be a considerable change in droplet transport and deposition associated with various compositions.

To unveil more underlying droplet transport dynamics that can be impacted by *V_d,i_* and θ, Figs. 7 (a)-(f) show the droplet residence time and droplet diameter change ratio (*d_d,f_*/*d_d,i_*) as a function of initial droplet diameter *d_d,i_*, spray cone angle θ, as well as the initial droplet composition. Specifically, Figs. 7 (a)-(c) show the droplet residence time with different initial droplet compositions (see Table 2 for details). It can be observed that the initial droplet composition has a negligible effect on droplet residence time, which is also due to the fact that the changes in the initial droplet composition do not significantly influence the initial droplet mass and evaporation/condensation characteristics. It can also be seen that droplet residence time is very short, indicating the quick deposition after injection as well as the short time for water evaporation or condensation. For example, vaccine droplets with *V_d,i_*=5 m/s need less than 0.03 seconds to reach OR. Accordingly, as shown in Figs. 7 (d)-(f), droplet size change was not significantly influenced by the initial droplet composition, *V_d,i_*, or θ. The evaporation can be considered the same for all cases, which is, as mentioned above, because they all consist of 95% evaporable water and 5% non-evaporable components. Therefore, water thermodynamic behavior is what determines droplet size change, which is the same for all cases. Droplet diameter change ratio *d_d,f_*/*d_d,i_* for different initial droplet compositions and θ are all higher than 99.5%. This is because of the short droplet residence time, which limited the total mass of water evaporation/condensation. In contrast, the droplet size change is noticeably influenced by the initial droplet diameter *d_d,i_*. Smaller *d_d,i_* induced higher water evaporation/condensation and droplet shrinkage, which is due to the Kelvin effect. Specifically, a smaller initial droplet diameter means higher surface curvature, leading to a faster evaporation/condensation rate.

**Fig. 7.**
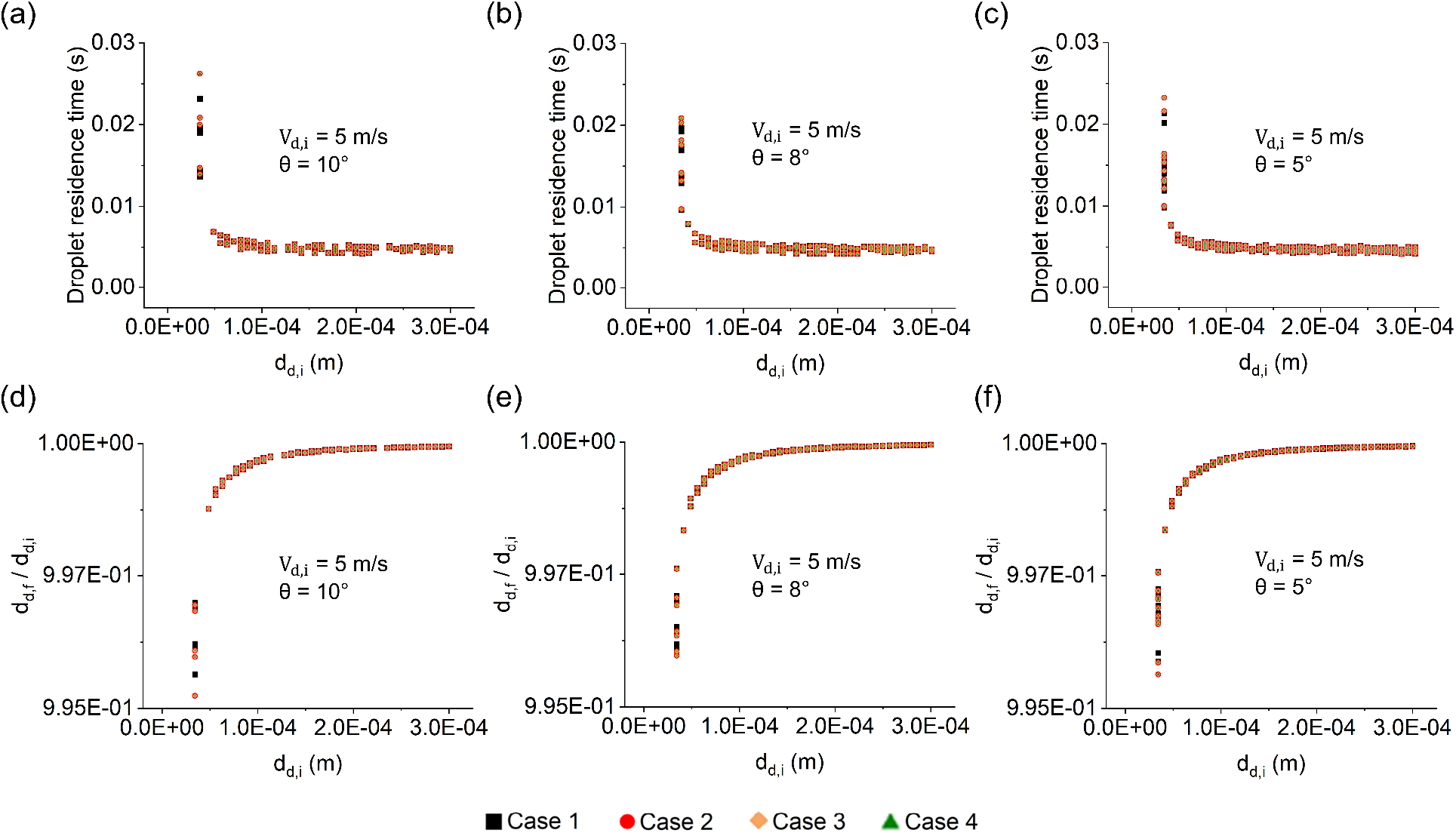
Vaccine droplet residence time and size change ratio as functions of d_d,i_, θ, and the initial droplet composition: (a-c) droplet residence time; and (d-f) droplet size change (*d_d,f_*/*d_d,i_*) versus droplet initial size *d_d,i_* for different spray cone angle θ with initial velocity *V_d,i_* of 5 m/s.

Figures 8 (a)-(c) plot the local distribution of vaccine droplets after they were sprayed into the nasal cavity. The effects of droplet compositions and cone angles on vaccine delivery to OR can be observed. It has been demonstrated that vaccine composition does not affect droplet distribution in OR/nasal cavity. The spray cone angle θ slightly affects droplet distribution. A wider cone angle (θ = 10°) leads to a relatively better VC of OR by droplets than a smaller cone angle (θ = 5°). Nonetheless, as it was discussed earlier, the simulation outcomes suggest that a larger cone angle θ can lead more droplets to be trapped in unexpected regions than the OR for SARS-CoV-2 vaccines. Assessment of VC and its influence on immune response and COVID-19 severity is a crucial step that needs to be taken to design an efficient intranasal vaccine spray.

**Fig. 8.**
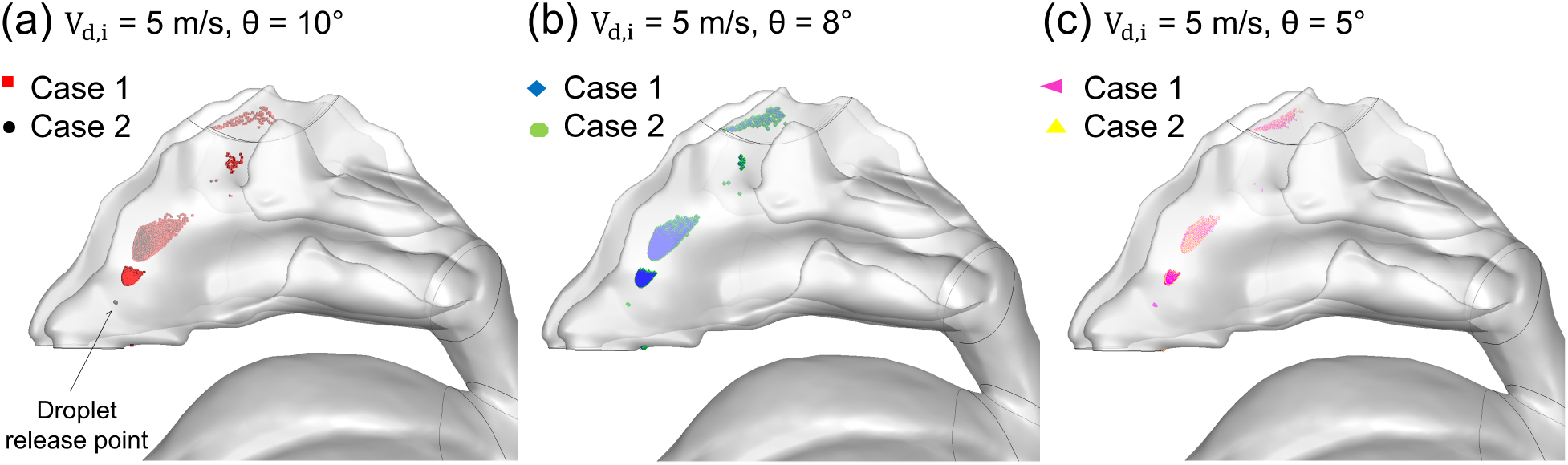
Local deposition of vaccine droplets in the nasal cavity for two initial droplet compositions (Case 1 and Case 2) at ***V_d,i_***=5 m/s with the spray cone angle (a) θ = 10°, (b) θ = 8°, and (c) θ = 5°.

### 6.3 Influence of nasal spray vaccine on viral replications predicted by the CFPD-HCD model

Figure 9 demonstrates the VC obtained from CFPD simulations and its influence on infection progression predicted by HCD analysis. In Fig. 9 (a), vaccine droplets (injected with different cone angle θ) deposited in OR are visualized. With θ = 15°, DF=1% of the droplets (see blue droplets in Fig. 9 (a)) can deposit in OR, which covers 1.8% of the OR surface area. Although the droplet deposition with θ = 15° is more scattered on the OR surface, the epithelial cells can be covered by those droplets are not higher than the rest due to the fact that fewer droplets are deposited. By reducing θ from 15° to 5°, higher DF (i.e., 5%) and VC (i.e., 4%) can be achieved. Nonetheless, reducing the cone angle further (i.e., 0° <θ < 5°) resulted in a decrease in VC despite the increased DF (see Fig. 9 (a)). Therefore, to seek the highest VC, θ =8° is recommended. In Fig. 9 (b), the impact of VC on viral titer kinetics is plotted. It can be observed that even with the highest VC, which is approximately 4% for θ = 8°, it is not able to trigger a sufficiently strong effect on varying the viral peak and its decaying trend. It is because that using the conventional atomization technique and available nasal spray nozzle openings, the vaccine droplets still cannot cover a sufficiently large area of OR (ACE-2 reach site). Therefore, the number of susceptible epithelial cells that can be infected by the virus for reproduction was not significantly reduced. However, if the vaccine droplet size distributions and release nozzle can be modified, it is possible that the VC can be further increased. Based on the comparisons shown in Fig. 9 (b), it can be hypothesized that increasing VC further can lead to a noticeable reduction in the viral titer peak, which has been proved by the comparisons shown in Fig. 10.

**Fig. 9.**
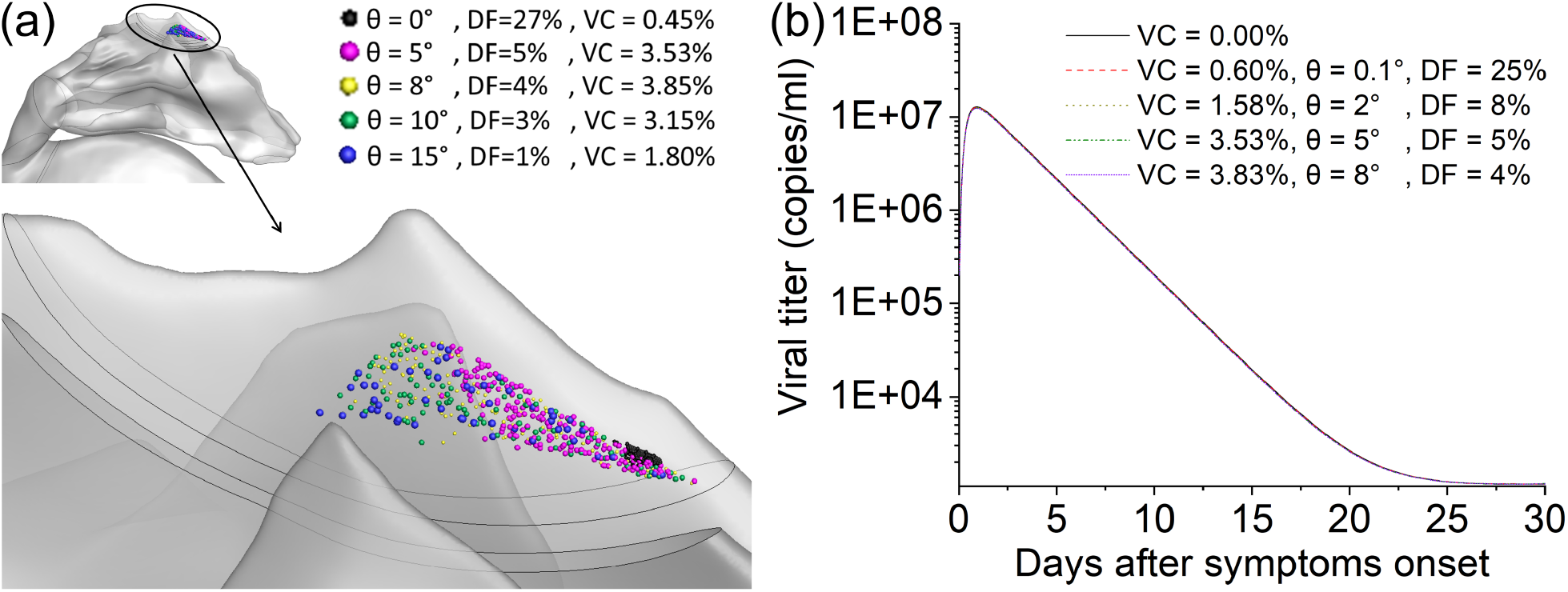
Vaccine coverage (VC) and its influence on viral titer kinetics predicted by the CFPD-HCD model: (a) Visualization of VC areas with different cone angles θ; (b) viral titer kinetics with different VCs obtained from CFPD simulation results.

**Fig. 10.**
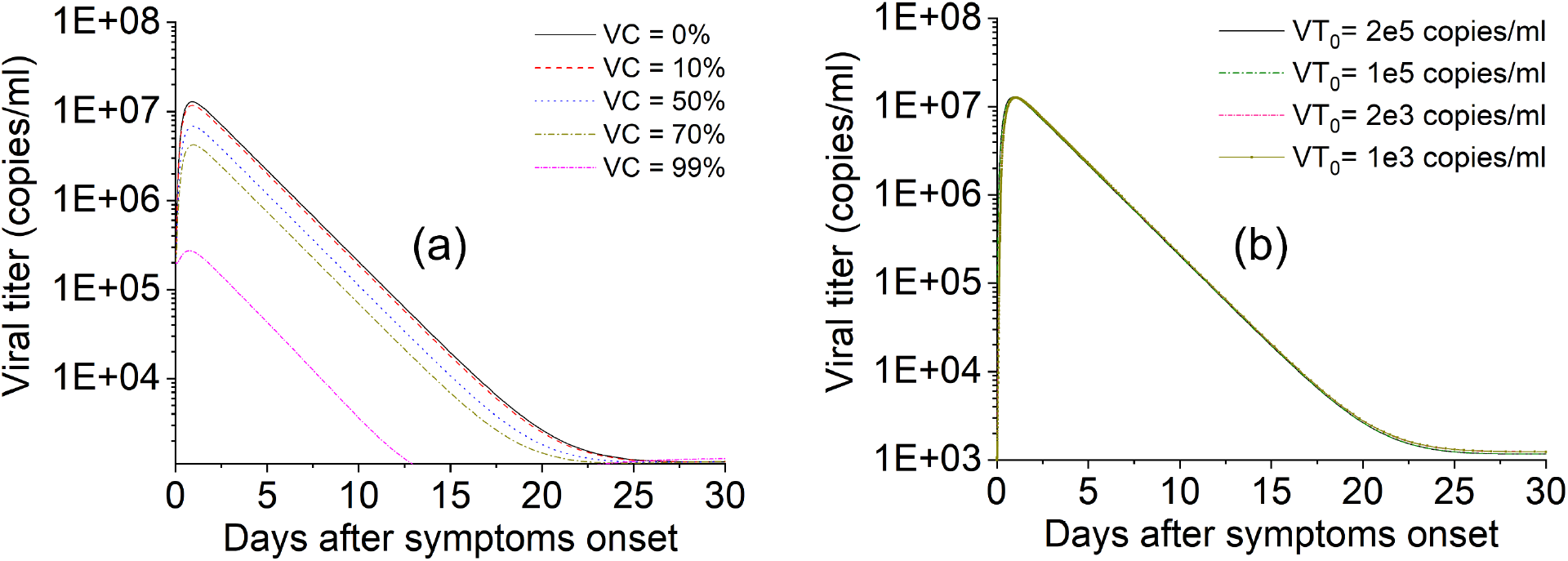
Effects of (a) vaccine coverage, and (b) initial viral titer on time profiles of SARS-CoV-2 viral titer.

### 6.4 HCD analysis of viral titer kinetics

To test the hypothesis mentioned at the end of Section 6.3, the effects of VC (the percentage of epithelial cells covered by the vaccine droplet deposition) and initial viral titer of SARS-CoV-2 on temporal dynamics of viral titer and its peak are predicted and compared in Fig. 10 (a). VC percentages were artificially varied from 0 % to 99%. Specifically, VC=0% represents the infection case without intranasal vaccine administration. It is shown that the increase in VC, which indicates fewer susceptible epithelial cells to be infected by SARS-CoV-2 invasion, can reduce the viral titer peak value. Indeed, the peak value of viral titer is reduced from 1e7 to 3e6 approximately if the vaccine droplets can cover up to 70% of the OR surface. It can also be found in Fig. 10 (a) that the viral agents will hardly duplicate themselves if VC can reach 99%. To further study the influence of the initial value of viral titer VT0 on the viral titer peak, VT0 was reduced, and the results are shown in Fig.10 (b) with VC=0%. It can be observed that changing from 1e3 copies/ml to 2e5 copies/ml, VT0 has negligible influence on the viral titer time profile if the numbers of susceptible epithelial cells are similar. Therefore, Figs. 10 (a) and (b) demonstrates that the effectiveness of the intranasal COVID-19 vaccine is highly dependent on the percentage of epithelial cells that can be covered by the aerosolized vaccine droplets. The nasal spray formulation and administration strategy must be optimized to maximize the VC, thereby reducing the possibility for SARS-CoV-2 reproduction.

## 10 Conclusions

Using an experimentally calibrated and validated CFPD-HCD model, this study predicted the transport of intranasal SARS-CoV-2 vaccine droplets delivery to ACE-2-rich regions (i.e., the olfactory region) in the nasal cavity of a 6-year-old female, as well as the vaccine influence on boosting local (mucosal) immunity against COVID-19 infection. Key conclusions are summarized below.

1. Vaccine droplet size change dynamics and deposition fractions in the olfactory region were not significantly influenced by initial droplet velocity or initial droplet composition.
2. Spray cone angle has a significant influence on vaccine droplet deposition fraction in the olfactory region, but has a negligible impact on droplet residence time and size change. To pursue the highest deposition fraction in the olfactory region, the most efficient cone angle is θ = 5°. In contrast, to cover the most epithelial cells in the olfactory region, the most efficient cone angle is θ = 8°.
3. Using the administration strategies investigated in this study, vaccine droplets can only occupy approximately 4% of the epithelial cells in the olfactory region, which is not sufficient for boosting immunity against SARS-CoV-2. To trigger effective immunity using intranasal COVID-19 vaccine spray, it is necessary to optimize the vaccine formulation and physical properties to enable the nasal spray to cover 99% of the epithelial cells in the olfactory region.

## 11 Limitations of the current study and future work

For this study, we encountered some limitations, and some simplifications needed to be made. The simplifications were:

1. Instead of a realistic breathing profile, an idealized sinusoidal waveform was employed for the CFPD simulation.
2. No mucus lining was included in the nasal cavity.
3. The subject was not systemically (intramuscularly) vaccinated.
4. The upper respiratory tract model was scaled down and connected to the larynx/trachea model of the 6-year-old child. Each vaccine droplet could cover the entire face cell it landed on. The lack of available clinical data for young children with COVID-19 caused us to use the adult clinical data that was reported in the literature.

In the future, to obtain more realistic flow field dynamics in the respiratory system, a realistic subject-specific breathing profile is recommended to be employed. Smaller spray droplets can be studied as well to see if they can improve vaccine delivery efficiency. Furthermore, it has been observed that mucus clearance and spray-liquid motion will alter the resting area of the spray droplets deposited on the mucus lining. Rygg, Hindle et al. (2016) showed that mucus clearance will transport the drugs that deposit on the mucus lining in the nasal cavity, which may or may not be in favor of drug delivery to targeted sites. Some of the drugs can be transported to the targeted site, some might be completely removed from the nasal cavity. Kolanjiyil, Alfaifi et al. (2022) reported that deposited spray liquid moved in the gravitational direction as well as inhalation flow direction using CFPD and Eulerian film model. Therefore, to provide more realistic resting site and VC of the COVID-19 vaccine droplets, post-deposition liquid film motion in the nasal cavity can be simulated by employing the Eulerian film model as a future work. Additionally, the entire upper respiratory system geometry of children could be constructed using CT/MRI images instead. To predict the local (mucosal) immune response triggered by nasal vaccines against COVID-19 more accurately using the HCD model, age-specific clinical data from young patients diagnosed with SARS-CoV-2 is recommended to be obtained for model calibration. Finally, the immunity triggered by SARS-CoV-2 intramuscular vaccines can be involved along with intranasal vaccines in HCD modeling to predict the robustness of topical and systemic immunization.

## Nomenclature

**Symbols**
*A_d_*: droplet surface area (m^2^)
*C_m_*: correction factor for Fuchs-Knudsen number
*C_p_*: specific heat of humid air (J/kg-K)
*C_p,d_*: specific heat of droplet (J/kg-K)
*D_w_*: water mass diffusivity (m^2^/s)
*d_d_*: droplet diameter (m)
*d_d,i_*: droplet initial diameter (m)
*d_d,f_*: droplet final diameter (m)
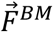: Brownian force (N)
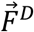: drag force (N)
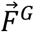: gravity (N)
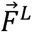: Saffman lift force (N)
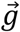: gravitational acceleration (m/s^2^)
*H_lat_*: latent heat (J/kg)
*K_w_*: Kelvin effect factor
*Kn*: Knudsen number
*k*: turbulence kinetic energy (J/kg)
*k_c_*: thermal conductivity (W/m-K)
*k_ct_*: turbulent thermal conductivity (W/m-K)
*k_hc_*: modified thermal conductivity (W/m^2^-K)
^*k_mc_*^: mass transfer coefficient (m/s)
*M_w_*: water molecular weight (kmol/kg)
*m_d_*: droplet mass (kg)
*Nu*: Nusselt number
*P_eq_*: equilibrium vapor pressure (Pa)
*P_sat_*: saturation vapor pressure (Pa)
*Pr*: Prandtl number
*R*: gas constant (J/mol-K)
*Re_d_*: droplet Reynolds number
*r_d_*: droplet radius (m)
*S_w_^m^*: mass source term (kg/m^3^-s)
*Sc*: Schmidt number
*Sh*: Sherwood number
*T*: humid airflow temperature (K)
*T_d_*: droplet temperature (K)
t: time (s)
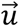: flow velocity (m/s)
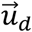: droplet velocity (m/s)
*V*: breathing velocity (m/s)
*V_d,i_*: droplet initial velocity (m/s)
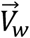: water molar volume (m^3^/kmol)
*y_w,surf_*: water mass fraction at droplet surface
*y_w,∞_*: water mass fraction in humid air mixture

**Greek symbols**
*α_m_*: mass accommodation factor
*λ*: gas mixture mean free path (m)
*μ*: viscosity of humid air (kg/m-s)
*μ _t_*: turbulent viscosity of humid air (kg/m-s)
*μ_d, surf_*: viscosity of humid air at droplet surface (kg/m-s)
*ρ*: density of humid airflow (kg/m^3^)
*ρ_w_*: density of water (kg/m^3^)
σ: droplet surface tension (N/m)
τ: breathing cycle time (s)
*Φ_exp_*: experimental values of reported for immune response
*Φ_HCD_*: computational values obtained for immune response
ω: specific rate of turbulence kinetic energy dissipation (J/kg-s)

**Acronyms**
*ACE-2*: angiotensin-converting enzyme 2
*CFPD*: computational fluid-particle dynamics
*COVID-19*: coronavirus disease 2019
*DF*: deposition fraction
*E_0_*: initial number of epithelial cells
*G*: generation
*GG*: gellan gum
*IgA*: immunoglobulin A
*IgG*: immunoglobulin G
*MRI*: magnetic resonance imaging
*N*: the number of fragments between max and min temperature
*NK*: natural killer cells
*ODEs*: ordinary differential equations
*OR*: olfactory region
*PBS*: phosphate-buffered saline
*RH*: relative humidity
*RMSE*: root mean squared error
*S*: spike protein
*SARS-CoV-2*: severe acute respiratory syndrome coronavirus 2
*SST*: shear stress transport
*TB*: tracheobronchial
*UDFs*: user-defined functions
*VC*: vaccine coverage
*VT_0_*: initial viral titer

## Acknowledgments

The research was made possible by funding through an award from the Oklahoma Center for the Advancement of Science and Technology (OCAST) (HR19-106). The research is also partially supported by the National Science Foundation (CBET 2120688) and the National Institutes of Health (NIH) Center of Biomedical Research Excellence (COBRE) (P20 GM103648). The use of Ansys software (Ansys Inc., Canonsburg, PA) as part of the Ansys-CBBL academic partnership coordinated by Dr. Thierry Marchal is gratefully acknowledged.

## Author contributions

Conceptualization, H.H., Y.F.; methodology, H.H., Y.F., X.C.; calibration and validation, H.H.; geometry preparation, E.K., C.F., Y.F., H.H.; simulations, H.H.; data curation, H.H.; data analysis, H.H.; writing-original draft, H.H.; writing-review and editing, H.H., E.K., X.C., Y.F.

## Disclaimer

It is not the intention of authors to provide specific medical advice but rather to provide readers with computational modeling details to better understand the fundamentals of fluid dynamics and host cell dynamics in COVID-19 nasal vaccine development. No *in vivo/in vitro* studies were conducted. Hence, no specific medical advice will be provided, and the authors urge you to consult with a qualified physician for answers to your personal medical concerns.

## Appendix

In Table A1, governing equations of the HCD model for non-specific immunity are provided. Viruses that bind to the susceptible and permissive target epithelial cells, *E*, infect them at a rate of *β_E_* VE. The infected cells are in the latent phase, *E_IL_*, at first and will become detectable, *E_I_*, at the rate of *δ_E_IL__E_IL_*. The infected cells die at the rate of *δ_E_I__E_I_*. Viruses use the machinery of the infected cells and replicate themselves at the rate of *π_V_E_I_*. Infectious virions that are released into the extracellular area of the lung tissue activate resting macrophages, M_R_, whose defensive mechanism is to phagocyte and dispose of dead cells and cell debris as well as invading influenza viruses, at the rate of *αVM_R_/V*_50_ + *V*. The death rate of tissue resting and activated macrophages are *δ_M_R__M_R_* and *δ_M_A__M_A_*, respectively. It has been reported that macrophages can become infected, M_I_, themselves with *λM_A_V* rate and replicate the viruses at the rate of *π_M_I__M_I_* (Pawelek, D. et al. 2016). Infected macrophages die at the rate of *δ_M_I__M_I_*.

Pre-death innate defensive response of the infected cells results in the production of interferons, F, by the rate of *π_F_E_I_*. Interferons send signals to the neighboring cells to be prepared to resist, *E_R_*, to a viral infection with ΦFE rate. The cells refracted to the infection will become susceptible to infection again at the rate of *δ_E_R__E_R_*. As well as signaling the neighboring uninfected cells, interferons activate resident natural killer, K, cells at the rate of *ϕ_k_F*K, which are lymphocytes of innate immunity, to secret cytokines and kill the detectable and latent-phased infected cells by inducing them to undergo apoptosis (Parham 1950) at the rate of *κ_k_E_I_*K and *κ_k_E_IL_*K, respectively. Resident and activated natural killer cells die respectively at the rate of *δ_k_K* and *δ_k_K_A_*. Resident dendritic cells, D, who act as sentinels take up pathogens and their products and become virus loaded, D_I_, by the rate of *β_D_DV*. The infected dendritic cells activate the natural killer cells to become effector cells. If effector natural killer cells outnumber the infected dendritic cells, they can kill the dendritic cells at the rate of δ_D_I__D_I_. However, if the natural killer cells are scarce and outnumbered by infected dendritic cells, they lead the infected dendritic cells to mature into the form that initiates adaptive immunity. Infected dendritic cells act as cellular messengers and migrate to the lymph node to call up adaptive immune responses (Parham 1950).

On encountering the antigen recognized by their antigen receptors in the lymphatic compartment, naïve CD4^+^ T cells and CD8^+^ T cells become activated by antigens at the rate of *π_H_* = *π*_*H*1_*D_M_*/(*π*_*H*2_ + *D_M_*) and *π_T_* = *π*_*T*1_*D_M_*/(*π*_*T*2_ + *D_M_*), respectively. Thereafter, activated CD4^+^ T and CD8^+^ T cells differentiate into effector CD4^+^ T cells (helper T cells) and CD8^+^ T cells (cytotoxic T cells) at the rate of *ρ_H_* = *ρ*_*H*1_*D_M_*/(*ρ*_*H*2_ + *D_M_*) and *ρ_T_* = *ρ*_*T*1_*D_M_*/(*ρ*_*T*2_ + *D_M_*), respectively. Through this process, effector CD4^+^ T cells help the mature dendritic cells to activate naïve CD8^+^ T cells. The death rate of effector CD4^+^ T and CD8^+^ T cells are δ_H_ = δ_H1_D_M_/δ_H2_ + D_M_ and δ_T_ = δ_T1_D_M_/δ_T2_ + D_M_, respectively. Thereafter, effector T cells travel to the infection site. Effector CD4^+^T cells are cytokines that only help cytotoxins to function. They do not directly eliminate infected cells. Cytotoxic T cells kill the infected cells at the rate of κ_E_E_I_γT_E_(t – τ_T_) during the adaptive immunity by inducing apoptosis, similar to natural killer cells during innate immunity. The model equations for adaptive immunity are provided in Table A2. Naïve B cells exposed to the antigens become activated at the rate of *π_B_* = *π*_*B*1_*D_M_*/(*π*_*B*2_ + *D_M_*) and next, they form a cognate pair with Helper CD4^+^ T cells to proliferate at the rate of *ρ_BA_* = *ρ*_*B*1_(*D_M_* + *hH_E_*)/(*D_M_* + *hH_E_* + *ρ*_*B*2_). Activated B cells differentiate into short-lived plasma cells and long-lived memory B cells at the rate of *π_S_B_A_* and π*_L_H_E_B_A_*, respectively. Short-lived plasma cells secret short-lived antibodies (IgM) at the rate of π*_AS_P_S_* whose clearance rate is δ*_AS_*A_*S*_. Long-lived plasma cells secret long-lived antibodies (IgG) at the rate of π*_AL_P_L_*. Long-lived antibodies have a clearance rate of δ*_AL_* A_*L*_. These antibodies circulate in the bloodstream and enter the infected site to neutralize viruses. Definition of model variables and their initial values are stated in Table A3.

The estimated values of parameters obtained from HCD model calibration with SARS-CoV-2 clinical data are presented in Table A4. To simplify the calibration, the conversion rate of cells that form latent phase to infected cells δ_EIL_= 3 day^-1^, death rate of infected epithelial cells δ_E_I__= 5 day^-1^, death rate of uninfected macrophages δ_MR_= 0.06 day^-1^, death rate of activated macrophages δ_M_A__ = 0.04 day^-1^, and the death rate of the infected macrophages δ_M_I__= 0.04 day^-1^ were kept constant (they are marked by a plus sign beside them in Table 4A). The values were chosen based on the data reported in the literature (Pawelek, D. et al. 2016, Vaidya, Bloomquist et al. 2021).

**Table A1.** ODEs of immune response in lung compartment

|  |  |
| --- | --- |
| $\frac{d}{dt} E = -\beta_E VE - \Phi FE + \delta_{E_R} E_R$ | (A1) |
| $\frac{d}{dt} E_{IL} = \beta_E VE - \delta_{E_{IL}} E_{IL} - \kappa_k E_{IL} K$ | (A2) |
| $\frac{d}{dt} E_I = \delta_{E_{IL}} E_{IL} - \delta_{E_I} E_I - \kappa_E E_I \gamma T_E(t - \tau_T) - \kappa_k E_I K$ | (A3) |
| $\frac{d}{dt} V = (1 - \varepsilon_1)(\pi_V E_I + \pi_{M_I} M_I) - c_V V - c_M M_A V - c_{A_S} A_S - c_{A_L} A_L$ | (A4) |
| $\frac{d}{dt} M_R = \delta_{M_R}(M_{R0} - M_R) - \frac{(1 - \varepsilon_2)\alpha VM_R}{V_{50} + V}$ | (A5) |
| $\frac{d}{dt} M_A = \frac{(1 - \varepsilon_2)\alpha VM_R}{V_{50} + V} - \lambda M_A V - \delta_{M_A} M_A$ | (A6) |
| $\frac{d}{dt} M_I = \lambda M_A V - \delta_{M_I} M_I$ | (A7) |
| $\frac{d}{dt} F = \pi_F E_I - \delta_F F$ | (A8) |
| $\frac{d}{dt} K = \delta_k(K_0 - K) - \phi_k FK$ | (A9) |
| $\frac{d}{dt} K_A = \phi_k FK - \delta_k K_A$ | (A10) |
| $\frac{d}{dt} E_R = \Phi FE - \delta_{E_R} E_R$ | (A11) |
| $\frac{d}{dt} D = \delta_D(D_0 - D) - \beta_D DV$ | (A12) |
| $\frac{d}{dt} D_I = \beta_D DV - \delta_{D_I} D_I$ | (A13) |

**Table A2.**
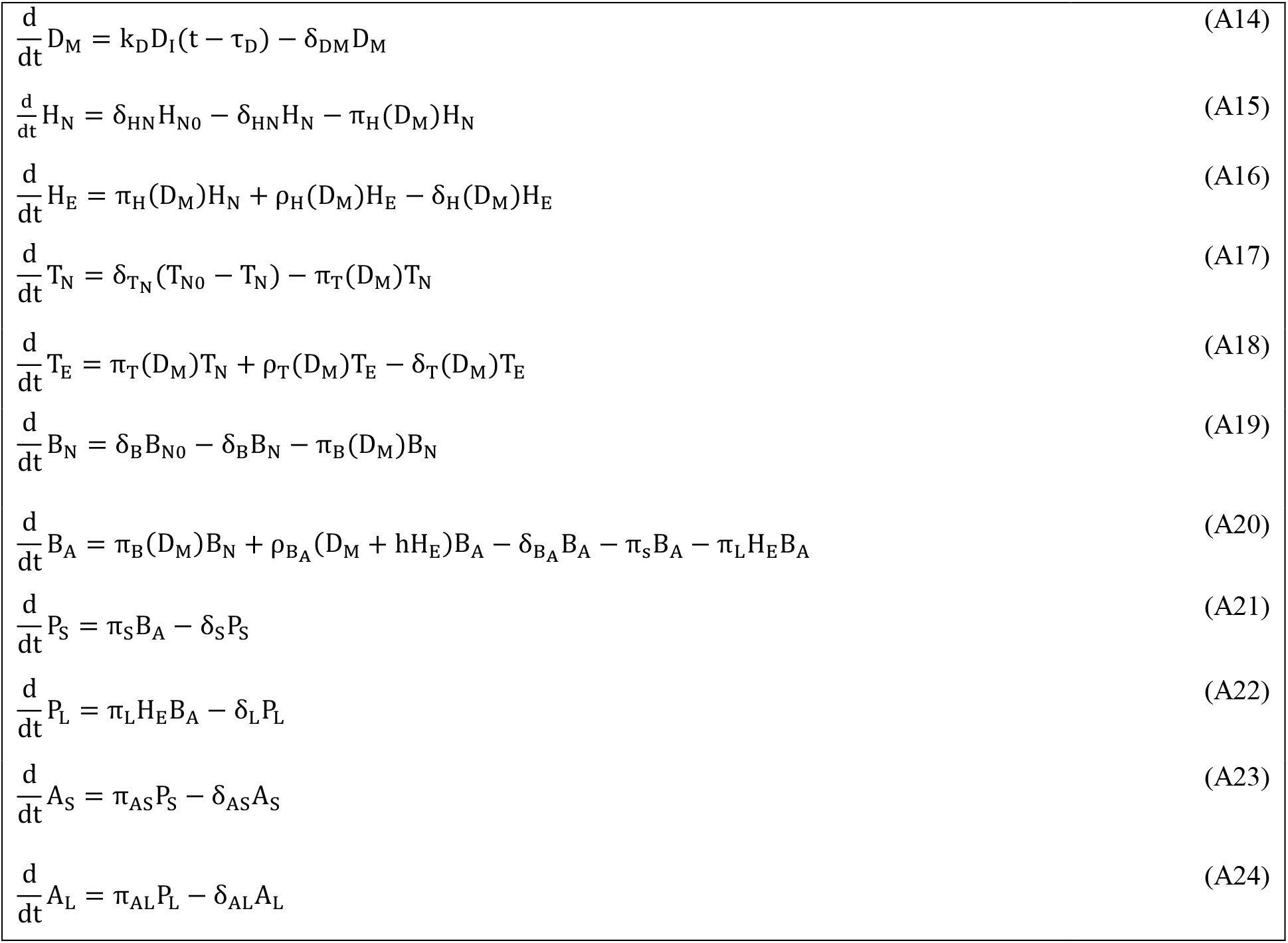
ODEs of immune response in lymphatic compartment

|  |  |
| --- | --- |
| $\frac{d}{dt}D_M = k_D D_I(t - \tau_D) - \delta_{DM} D_M$ | (A14) |
| $\frac{d}{dt}H_N = \delta_{HN} H_{N0} - \delta_{HN} H_N - \pi_H(D_M) H_N$ | (A15) |
| $\frac{d}{dt}H_E = \pi_H(D_M) H_N + \rho_H(D_M) H_E - \delta_H(D_M) H_E$ | (A16) |
| $\frac{d}{dt}T_N = \delta_{TN}(T_{N0} - T_N) - \pi_T(D_M) T_N$ | (A17) |
| $\frac{d}{dt}T_E = \pi_T(D_M) T_N + \rho_T(D_M) T_E - \delta_T(D_M) T_E$ | (A18) |
| $\frac{d}{dt}B_N = \delta_B B_{N0} - \delta_B B_N - \pi_B(D_M) B_N$ | (A19) |
| $\frac{d}{dt}B_A = \pi_B(D_M) B_N + \rho_{BA}(D_M + h H_E) B_A - \delta_{BA} B_A - \pi_S B_A - \pi_L H_E B_A$ | (A20) |
| $\frac{d}{dt}P_S = \pi_S B_A - \delta_S P_S$ | (A21) |
| $\frac{d}{dt}P_L = \pi_L H_E B_A - \delta_L P_L$ | (A22) |
| $\frac{d}{dt}A_S = \pi_{AS} P_S - \delta_{AS} A_S$ | (A23) |
| $\frac{d}{dt}A_L = \pi_{AL} P_L - \delta_{AL} A_L$ | (A24) |

**Table A3.** Definition of HCD variables and their initial values

| Variable | Definition | Initial value |
| --- | --- | --- |
| E | Uninfected epithelial cells | 4e7 |
| E<sub>IL</sub> | Epithelial cells in eclipse phase | 0 |
| E<sub>I</sub> | Infected epithelial cells | 0 |
| V | Viral load (titer) | 2e5 |
| M<sub>R</sub> | Uninfected resting macrophages circulating within the host | 1e3 |
| M<sub>A</sub> | Activated macrophages at the site of the infection | 0 |
| M<sub>I</sub> | Productively infected macrophages | 0 |
| F | Interferons | 0 |
| K | Resident NK cells | 1e3 |
| K<sub>A</sub> | Activated NK cells | 0 |
| E<sub>R</sub> | Uninfected cells that are refractory to infection | 0 |
| D | Immature dendritic cells | 1e3 |
| D<sub>I</sub> | Virus-loaded dendritic cells | 0 |
| D<sub>M</sub> | Mature dendritic cells | 0 |
| H<sub>N</sub> | Naïve CD4 T cells | 1e3 |
| H<sub>E</sub> | Helper T cell | 0 |
| T<sub>N</sub> | Naïve CD8 T cells | 1e3 |
| T<sub>E</sub> | Effector CD8 T cells | 0 |
| B<sub>N</sub> | Naïve B cells | 1e3 |
| B<sub>A</sub> | Activated B cells | 0 |
| P<sub>S</sub> | Short-lived antibody-secreting plasma cells | 0 |
| P<sub>L</sub> | Long-lived antibody-secreting cells | 0 |
| A<sub>S</sub> | Short-lived antibody | 0 |
| A<sub>L</sub> | Long-lived antibody | 0 |

**Table A4.** Definitions and values of coefficients in the HCD model

| Parameter | Definition | Value (bounds) |
| --- | --- | --- |
| $\beta_E$ | Infection rate of uninfected epithelial cells ($\text{day}^{-1} (\text{copies/ml})^{-1}$) | (1e-8,5.5e-4)<br>3e-4 |
| $\Phi$ | Interferon-induced antiviral efficacy ((IFN fold change) $\text{day}^{-1}$) | (1e-8,1e-4)<br>1.1e-5 |
| $\delta_{ER}$ | Reversion rate from refractory ($\text{day}^{-1}$) | (2,5)<br>2.97 |
| $\delta_{EIL}$ | Conversion rate of cells form latent phase to infected ($\text{day}^{-1}$) | 3<sup>+</sup> |
| $\kappa_k$ | Killing rate of infected cells by NK cells ($\text{day}^{-1}$) | (1e-4,1e-2)<br>5.2e-3 |
| $\delta_{EI}$ | Death rate of infected epithelial cells ($\text{day}^{-1}$) | 5<sup>+</sup> |
| $\kappa_E$ | Killing rate of infected cells by cytotoxic ($\text{CD8}^+$) T effectors ($\text{day}^{-1}$) | (1e-5,1e-3)<br>8.4e-4 |
| $\varepsilon_1$ | The drug efficacy | 0 |
| $\pi_V$ | Virus production rate per infected epithelial cells ( ($\text{copies/ml}$) $\text{day}^{-1}$) | (2,6)<br>2.61 |
| $\pi_{M_I}$ | Virus production rate per infected macrophages ( ($\text{copies/ml}$) $\text{day}^{-1}$) | (0.01,0.08)<br>0.07 |
| $c_V$ | Virus nonspecific clearance rate ($\text{day}^{-1}$) | (0.1,5)<br>0.5 |
| $c_M$ | Virus clearance rate through death of infected macrophages ($\text{day}^{-1}$) | (1e-4,1e-2)<br>4.3e-3 |
| $c_{AS}$ | Rate of virus neutralization by short-lived antibodies ($\text{day}^{-1} \mu\text{g}^{-1}$) | (0.05,2)<br>5.3e-1 |
| $c_{AL}$ | Rate of virus neutralization by long-lived antibodies ($\text{day}^{-1} \mu\text{g}^{-1}$) | (0.05,2)<br>1.1 |
| $\delta_{MR}$ | Death rate of uninfected macrophages ($\text{day}^{-1}$) | 0.06<sup>+</sup> |
| $\varepsilon_2$ | The effect of an anti-inflammatory drug efficacy | 0 |
| $\alpha$ | Activation rate of macrophages ($\text{day}^{-1}$) | (0.01,0.02)<br>1.8e-2 |
| $V_{50}$ | Viral load for half-maximal activation of macrophages | (2e-4,3e-4)<br>2.9e-4 |
| $\lambda$ | Infection rate of the activated macrophages | (1e-4,1e-2)<br>4.4e-4 |

**Table A4.** Definitions and values of coefficients in the HCD model - continued
| Parameter | Definition | Value (bounds) |
| --- | --- | --- |
| $\delta_{MA}$ | Death rate of activated macrophages ($\text{day}^{-1}$) | 0.04<sup>+</sup> |
| $\delta_{MI}$ | Death rate of the infected macrophages ($\text{day}^{-1}$) | 0.04<sup>+</sup> |
| $\pi_F$ | Secretion rate of interferons by infected cells ((IFN fold change) $\text{day}^{-1} \text{ cell}^{-1}$) | (1e-5,1e-3)<br>0.9e-3 |
| $\delta_F$ | Decay rate of interferons ($\text{day}^{-1}$) | (3,7)<br>5 |
| $\delta_k$ | Death rate of NK cells ($\text{day}^{-1}$) | (2e-5,4e-5)<br>3.5e-5 |
| $\phi_k$ | Activation rate of NK cells by interferons | (1e-5,1e-3)<br>0.4e-3 |
| $\delta_D$ | Death rate of immature dendritic cells ($\text{day}^{-1}$) | (9e-4,1e-3)<br>0.97e-3 |
| $\beta_D$ | Infection rate of dendritic cells by unit Corona virus ($\text{day}^{-1} (\text{EID}_{50}/\text{ml})^{-1}$) | (1e-3,3e-3)<br>2.3e-3 |
| $\delta_{DI}$ | Death rate of infected DCs ($\text{day}^{-1}$) | (5.5,6.5)<br>6.2 |
| $k_D$ | Antigen processing rate ($\text{day}^{-1}$) | (2,4)<br>2.5 |
| $\delta_{HN}$ | Death rate of naïve $\text{CD8}^+$ T cells ($\text{day}^{-1}$) | (7e-6,8e-6)<br>7.9e-6 |
| $\pi_{H1}$ | Maximum activation rate of naïve $\text{CD4}^+$ T cells ($\text{day}^{-1}$) | (1.2,1.4)<br>1.26 |
| $\pi_{H2}$ | Number of $D_M$ cells for half-maximal activation of effector $\text{CD4}^+$ T cells | (120,140)<br>133 |
| $\rho_{H1}$ | Maximum proliferation rate of naïve $\text{CD4}^+$ T cells ($\text{day}^{-1}$) | (0.1-0.2)<br>0.17 |
| $\rho_{H2}$ | Number of $D_M$ cells for half-maximal proliferation of effector $\text{CD4}^+$ T cells | (3e3-4e3)<br>3.15e+3 |
| $\delta_{H1}$ | Maximum clearance rate of effector $\text{CD4}^+$ T cells ($\text{day}^{-1}$) | (2-2.5)<br>2.4 |
| $\delta_{H2}$ | Number of $D_M$ cells for half-maximal clearance of effector $\text{CD4}^+$ T cells | (1.4,1.5)<br>1.43 |
| $\delta_{TN}$ | Death rate of naïve $\text{CD8}^+$ T cells ($\text{day}^{-1}$) | (6e-3-7e-3)<br>6.3e-3 |

**Table A4.** Definitions and values of coefficients in the HCD model - continued
| Parameter | Definition | Value (bounds) |
| --- | --- | --- |
| $\pi_{T1}$ | Maximum activation rate of CD8<sup>+</sup> T cells from naïve CD8<sup>+</sup> T cells (day<sup>-1</sup>) | (2,4)<br>2.3 |
| $\pi_{T2}$ | Number of D<sub>M</sub> cells for half maximal activation of naïve CD8<sup>+</sup> T cells | (25,35)<br>25.2 |
| $\rho_{T1}$ | Maximum proliferation rate of naïve CD8<sup>+</sup> T cells (day<sup>-1</sup>) | (1.7,1.9)<br>1.8 |
| $\rho_{T2}$ | Number of D<sub>M</sub> cells for half-maximal proliferation of effector CD8<sup>+</sup> T cells | (300,350)<br>325 |
| $\delta_{T1}$ | Maximum clearance rate of effector CD8<sup>+</sup> T cells (day<sup>-1</sup>) | (0.5,0.7)<br>0.68 |
| $\delta_{T2}$ | Number of D<sub>M</sub> cells for half-maximal clearance of effector CD8<sup>+</sup> T cells | (3.5,5.5)<br>3.6 |
| $\pi_{B1}$ | Maximum activation rate of naïve B cells (day<sup>-1</sup>) | (2.5,3.5)<br>2.9 |
| $\pi_{B2}$ | Number of D<sub>M</sub> cells for half-maximal activation of naïve B cells | (1e4,2.5e4)<br>1.2e-4 |
| $\delta_B$ | Death rate of naïve B cells (day<sup>-1</sup>) | (2e-3,3e-3)<br>2.5e-3 |
| $\rho_{B1}$ | Maximum proliferation rate of activated B cells (day<sup>-1</sup>) | (0.9,1.1)<br>0.92 |
| $\rho_{B2}$ | Number of D<sub>M</sub> cells for half-maximal proliferation of activated B cells | (4e3,5e3)<br>4.8e3 |
| h | Factor of B-cell activation by CD4<sup>+</sup> T cells | (0.7,0.9)<br>0.75 |
| $\delta_{BA}$ | Clearance rate of activated B cells (day<sup>-1</sup>) | (0.4,0.6)<br>0.60 |
| $\pi_s$ | Differentiation rate of activated B cells into short-lived antibody-secreting plasma cells (day<sup>-1</sup>) | (0.4,0.6)<br>0.51 |
| $\pi_L$ | Differentiation rate of activated B cells into long-lived antibody-secreting plasma cells (day<sup>-1</sup>) | (0.4,0.6)<br>0.44 |

**Table A4.** Definitions and values of coefficients in the HCD model - Continued
| Parameter | Definition | Value (bounds) |
| --- | --- | --- |
| $\delta_S$ | Death rate of short-lived antibody-secreting plasma cells ($\text{day}^{-1}$) | (0.01,0.5)<br>0.05 |
| $\delta_L$ | Death rate of long-lived antibody-secreting plasma cells ($\text{day}^{-1}$) | (0.02,0.03)<br>0.28e-1 |
| $\pi_{AS}$ | Secretion rate of antibody titer by unit short-lived plasma call ($\text{day}^{-1}$) | (0.3,0.8)<br>0.67 |
| $\delta_{AS}$ | Clearance rate of short-lived antibody ($\text{day}^{-1}$) | (0.05,0.15)<br>0.14 |
| $\pi_{AL}$ | Secretion rate of antibody titer by unit long-lived plasma call ($\text{day}^{-1}$) | (0.3,0.8)<br>0.43 |
| $\delta_{AL}$ | Clearance rate of long-lived antibody ($\text{day}^{-1}$) | (0.05,0.15)<br>0.13 |

